# Feeding state functionally reconfigures a sensory circuit to drive thermosensory behavioral plasticity

**DOI:** 10.1101/2020.06.22.164152

**Authors:** Asuka Takeishi, Jihye Yeon, Nathan Harris, Wenxing Yang, Piali Sengupta

**Affiliations:** Department of Biology, Brandeis University, Waltham, MA 02454; Neural circuit of multisensory integration RIKEN Hakubi Research Team, RIKEN Cluster for Pioneering Research, RIKEN Center for Brain Science, Wako, Japan; Department of Organismic and Evolutionary Biology, Center for Brain Science, Harvard University, Cambridge, MA 02138; Department of Physiology, West China School of Basic Medical Sciences and Forensic Medicine, Sichuan University, Chengdu, Sichuan, China

**Author notes:** These authors contributed equally to this work (Asuka Takeishi, Jihye Yeon).

## Abstract

Internal state alters sensory behaviors to optimize survival strategies. The neuronal mechanisms underlying hunger-dependent behavioral plasticity are not fully characterized. Here we show that feeding state regulates *C. elegans* negative thermotaxis behavior by engaging a modulatory circuit whose activity gates the output of the core thermotaxis network. Feeding state does not alter the activity of the core thermotaxis circuit comprised of AFD thermosensory and AIY interneurons. Instead, prolonged food deprivation potentiates temperature responses in the AWC sensory neurons, which inhibit the postsynaptic AIA interneurons to override and disrupt AFD-driven thermotaxis behavior. Acute inhibition and activation of AWC and AIA, respectively, restores negative thermotaxis in starved animals. We find that state-dependent modulation of AWC-AIA temperature responses requires INS-1 insulin-like peptide signaling from the gut and DAF-16 FOXO function in AWC. Our results describe a mechanism by which functional reconfiguration of a sensory network via gut-brain signaling drives state-dependent behavioral flexibility.

## Introduction

Responses of animals to sensory stimuli are extensively modulated by their internal state (Grunwald Kadow, 2019; Kim et al., 2017; Li and Dulac, 2018; Stowers and Liberles, 2016). The sex and hormonal conditions of an animal determine its responses to pheromones (Li and Dulac, 2018; Martin-Sanchez et al., 2015; McGrath and Ruvinsky, 2019; Stowers and Liberles, 2016), and behavioral arousal thresholds are regulated by sleep-wake cycles (Allada et al., 2017; Lee and Dan, 2012). A particularly well-studied internal state is that of satiety. Well-fed animals exhibit distinct responses to environmental cues compared to animals that have been food-deprived (Augustine et al., 2020; Kim et al., 2017). Starvation not only modulates responses to food-related chemical cues, but also generally and broadly regulates animal behaviors (Dietrich et al., 2015; Rengarajan et al., 2019; Sayin et al., 2019; Trent et al., 1983; Yang et al., 2015). These behavioral changes may allow animals to prioritize food seeking over other behavioral drives. How starvation signals are integrated to alter neuron and circuit properties are not fully understood.

Neuromodulation is a major mechanism driving satiety state-dependent behavioral plasticity. Modulation of interoceptive hypothalamic circuits by circulating hormones regulates appetitive behaviors in satiated and starved animals (Andermann and Lowell, 2017; Augustine et al., 2020). In food-deprived *Drosophila*, increased attraction and decreased repulsion to appetitive and aversive stimuli, respectively, are mediated via parallel modulation of attractive and aversive chemosensory circuits by diverse neuromodulators [eg. (Inagaki et al., 2014; Ko et al., 2015; Marella et al., 2012; Root et al., 2011; Vogt et al., 2020)]. Prior food deprivation or pairing starvation with a stimulus also markedly alters sensory responses in *C. elegans* via monoaminergic and neuropeptidergic signaling among others [eg.(Chalasani et al., 2010; Cho et al., 2016; Ezcurra et al., 2011; Ghosh et al., 2016; Rengarajan et al., 2019; Saeki et al., 2001; Tomioka et al., 2006; Torayama et al., 2007)]. In addition to targeting central interoceptive circuits, these neuromodulators can also mediate presynaptic facilitation or inhibition at the first sensory synapse (Devineni et al., 2019; Inagaki et al., 2014; Ko et al., 2015; Rengarajan et al., 2019; Root et al., 2011), or directly tune sensory responses (Cho et al., 2016; Ezcurra et al., 2011; Oda et al., 2011). It is currently unclear how generalizable these principles are, and whether alternate mechanisms also contribute to the generation of feeding state-dependent behavioral plasticity.

Thermotaxis navigation behaviors in *C. elegans* are particularly susceptible to feeding state. When placed on a spatial thermal gradient, well-fed but not food-deprived *C. elegans* navigates towards temperatures at which they were cultivated (*T*_*c*_) for 3-4h prior to the behavioral assay (Figure 1A) (Hedgecock and Russell, 1975). Thermosensation is mediated primarily by the AFD sensory neurons, and their major postsynaptic partners, the AIY interneurons in the head of *C. elegans* (Mori and Ohshima, 1995). Additional sensory neurons including the AWC olfactory neurons also exhibit temperature responses but play relatively minor roles in regulating thermotaxis behaviors under standard assay conditions (Biron et al., 2008; Ikeda et al., 2020; Kuhara et al., 2008). Temperature responses in AFD appear to be indifferent to feeding state (Biron et al., 2006; Ramot et al., 2008a; Tsukada et al., 2016), leaving open the question of which circuit mechanisms integrate internal state information into the thermotaxis circuit to disrupt thermotaxis.

**Figure 1.**
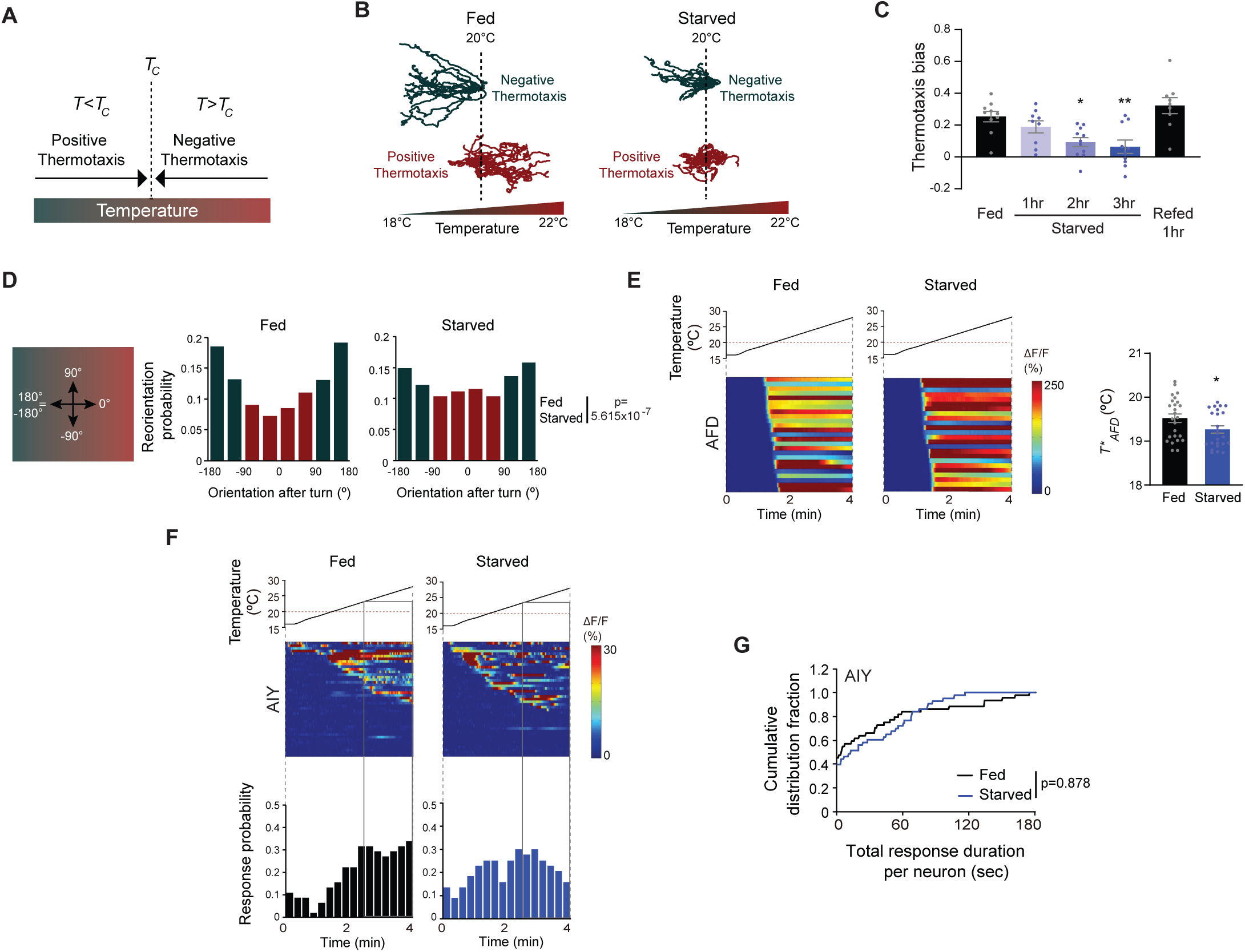
Starvation disrupts negative thermotaxis but does not affect temperature responses in AFD and AIY. **A)** Schematic of experience-dependent thermotaxis behavior of *C. elegans* (Hedgecock and Russell, 1975). *T*: ambient temperature on gradient; *T*_*c*_ = cultivation temperature 3-4h prior to assay. Warm and cool temperatures are indicated in red and green, respectively. **B)** Tracks of individual worms on a long linear thermal gradient from a single representative assay of ∼15 animals each. Worms were cultivated at 15°C or 25°C for negative (green tracks) or positive (red tracks) thermotaxis assays, respectively, with or without bacterial food for 3h prior to assay. Dashed lines indicate the temperature (20°C) at which animals were placed at the start of the assay. Tracks were superimposed post analysis for presentation. **C)** Mean thermotaxis bias of animals subjected to the indicated feeding conditions on a short thermal gradient. Thermotaxis bias was calculated as [(run duration toward colder side – run duration toward warmer side)/total run duration]. Each dot represents the thermotaxis bias of a biologically independent assay comprised of 15 animals. Errors are SEM. * and ** indicates different from fed at p<0.05 and p<0.01, respectively (ANOVA with Tukey’s multiple comparisons test). **D)** (Left) Schematic of track orientation on a linear thermal gradient. Orientation parallel to the gradient towards warm temperatures is 0°, orientation orthogonal to the gradient is 90° or -90°, and orientation parallel to the gradient towards cold temperatures is 180° or-180°. (Right) Histograms of movement orientation following a turn. Tracks from 8 assays of 15 animals each were categorized into bins of 45°. Red and green bars indicate orientation toward the warmer/orthogonal or cooler side, respectively. The p-value was derived using the Mardia-Watson-Wheeler non-parametric test for circular data. **E)** (Left) Intracellular calcium dynamics in AFD expressing GCaMP6s in response to a linear rising temperature stimulus (black lines) at 0.05°C/sec. Red dashed line indicates cultivation temperature of 20°C. Each row in the heatmaps displays responses from a single AFD neuron ordered by the time of the first response; n =25 (fed) and 24 (starved). (Right) Mean *T\**_*AFD*_ of fed and starved animals calculated from data shown in heatmaps at left. Each dot is the *T\**_*AFD*_ of a single neuron. Errors are SEM. * indicates different from fed at p<0.05 (Student’s t-test). **F)** (Top) Intracellular calcium dynamics in AIY expressing GCaMP6s in response to a linear rising temperature stimulus (black lines) at 0.05°C/sec. Red dashed line indicates cultivation temperature of 20°C. Each row in the heatmaps displays responses from a single neuron ordered by the time of the first response; n = 44 (fed) and 43 (starved). (Bottom) Each bar in the histograms represents the proportion of neurons responding during 15s bins. The behavioral temperature range of 23°C-28°C is indicated by vertical solid gray lines. **G)** Cumulative distribution fraction plots of total duration of calcium responses per AIY neuron calculated from data shown in **F**. Distributions were compared using the Kolmogorov-Smirnov test. Also see Figure S1.

A previous study implicated insulin signaling in the regulation of feeding state-dependent thermotaxis behavioral plasticity in *C. elegans* (Kodama et al., 2006). The INS-1 insulin-like peptide (ILP) gene was suggested to antagonize the DAF-2 insulin receptor and the DAF-16/FOXO transcription factor to disrupt thermotaxis behavior upon food deprivation, such that starved *ins-1* mutants continue to perform thermotaxis (Kodama et al., 2006). However, neither the source of INS-1 production, nor its site of action were definitively identified. Behavioral experiments suggested that INS-1 expression from subsets of neurons target the AIY, AIZ and RIA interneurons implicated in the thermotaxis circuit (Kodama et al., 2006), and temperature responses in AIZ were shown to be regulated as a function of satiety state (Kodama et al., 2006). However, given conflicting reports on the roles of AIZ and RIA in driving thermotaxis behaviors (Luo et al., 2014a; Mori and Ohshima, 1995; Ohnishi et al., 2011), how internal feeding state and ILP signaling modulate the thermotaxis circuit to alter navigation behaviors remain unclear.

Here we show that internal feeding state regulates thermotaxis behavioral plasticity via INS-1-mediated neuromodulation of the AWC sensory and postsynaptic AIA interneurons. We find that although temperature responses in neither AFD nor AIY are altered upon prolonged food deprivation, the probability and duration of temperature responses in AWC are increased under these conditions. AWC inhibits the AIA interneurons via glutamatergic signaling to alter locomotory strategies and disrupt AFD-driven thermotactic navigation in starved animals. We show that expression of *ins-1* specifically in the gut is necessary for internal state-dependent thermotaxis behavioral plasticity, and establish that gut-derived INS-1 signaling targets DAF-16/FOXO in AWC to regulate temperature responses and circuit activity in response to feeding state. Our results indicate that internal state drives thermosensory behavioral plasticity by tuning the activity state of a modulatory circuit via gut-to-brain signaling. In turn, this circuit gates the ability of the core thermotaxis network to regulate navigational strategies as a function of environmental temperature changes.

## Results

### Prolonged food deprivation disrupts thermotaxis navigation behavior

When placed at temperatures (*T*) warmer than the *T*_*c*_, *C. elegans* moves towards cooler temperatures (negative thermotaxis) (Hedgecock and Russell, 1975) (Figure 1A). Conversely, when placed at *T<T*_*c*_, animals move towards warmer temperatures (positive thermotaxis) (Figure 1A). To characterize the effects of prolonged starvation on thermotaxis at high resolution, we examined animal movement under assay conditions that permitted both negative and positive thermotaxis (Luo et al., 2014a). Well-fed young adult hermaphrodites grown at 15°C, and placed at 20°C at the center of a shallow linear thermal gradient (long thermal gradient; see Methods), moved robustly down the gradient towards cooler temperatures (Figure 1B). Conversely, animals grown at 25°C and placed at 20°C on this gradient moved towards warmer temperatures (Figure 1B). Food deprivation for 3h disrupted both navigation behaviors (Figure 1B) (Chi et al., 2007; Ramot et al., 2008b). In particular, examination of individual animal trajectories showed that food-deprived animals grown at 15°C exhibited more sharp turns and reversals than their fed counterparts, resulting in prolonged residence at the starting temperature. While a subset of food-deprived 15°C-grown animals eventually moved down the gradient, starved 25°C-grown animals essentially remained at their starting temperature throughout the assay (Figure 1B).

While negative thermotaxis is exhibited across a range of assay parameters, positive thermotaxis is typically consistently observed only under a relatively restricted set of assay conditions (Jurado et al., 2010; Ramot et al., 2008b). We chose to further pursue the effects of food deprivation on negative thermotaxis which in addition to being robust, can also be performed at higher throughput (Clark et al., 2007; Ryu and Samuel, 2002). On spatial thermal gradients, negative thermotaxis is mediated primarily via klinokinesis (Clark et al., 2007; Luo et al., 2014a; Ryu and Samuel, 2002; Zariwala et al., 2003). In this behavioral strategy, worms moving towards cooler temperatures suppress reorientations (turns) and consequently, extend the duration of forward movement (runs). Conversely, when moving towards the non-preferred warmer temperatures, worms increase turn frequency and decrease run duration. In parallel, following a turn, worms preferentially bias the direction of a new run towards cooler temperatures (Luo et al., 2014a). These strategies result in net migration of animals down the gradient. We asked whether either or both strategies are disrupted upon starvation.

To quantify klinokinesis, we calculated thermotaxis bias [(run duration towards cold – run duration towards warm)/ total run duration] of adult *C. elegans* hermaphrodites grown at *T*_*c*_=20°C and navigating a steeper linear thermal gradient (short thermal gradient; see Methods) (Chi et al., 2007; Clark et al., 2007). While well-fed animals exhibited robust negative thermotaxis bias under these conditions, animals starved for >2h were essentially athermotactic, indicating that these animals were unable to modulate turning frequency as a function of experienced temperature changes (Figure 1C) (Chi et al., 2007; Hedgecock and Russell, 1975; Kodama et al., 2006). We established that starved animals also failed to orient run direction towards cooler temperatures following a turn, and instead oriented their runs near-randomly on the gradient (Figure 1D). Average velocities of fed and starved animals were indistinguishable on these gradients (average velocity: fed – 117 μm/s, starved for 3h – 106 μm/s; n=10 animals each). Together, these observations indicate that prolonged starvation abolishes both klinokinesis and biased run direction to disrupt negative thermotaxis. All subsequent experiments using starved animals were performed following food removal for 3h (referred to interchangeably as starvation or food deprivation).

We tested whether starvation-mediated alteration of negative thermotaxis is reversible. Refeeding starved animals for an hour was sufficient to restore robust negative thermotaxis bias (Figure 1C) (Chi et al., 2007; Mohri et al., 2005), indicating that prolonged starvation does not irreversibly alter the function of the underlying circuit. Since starvation deprives animals of chemosensory inputs from bacteria and alters internal metabolic state, we asked whether exposure to bacterial odors was sufficient to mimic the fed state. However, the presence of bacteria on the lids of agar plates did not override the effect of starvation on negative thermotaxis (Figure S1A). Moreover, feeding animals either live or antibiotic-killed bacteria was sufficient to mimic the well-fed condition for negative thermotaxis (Figure S1B). We infer that prolonged starvation alters internal state to disrupt negative thermotaxis.

### Starvation does not alter AFD or AIY temperature responses

The bilateral pair of AFD sensory neurons are the primary thermoreceptors driving thermotaxis navigation behaviors (Goodman and Sengupta, 2018; Mori and Ohshima, 1995). To assess whether AFD temperature responses are modulated by food, we examined calcium dynamics in fed and starved animals grown at 20°C expressing GCaMP6s specifically in AFD and subjected to a rising temperature stimulus. To more closely mimic the temperature changes that animals experience when they are navigating the linear thermal gradient used in our assays, we performed all measurements using a shallow (0.05°C/sec) linear rising temperature ramp from 16°C-28°C (see Methods). AFD responds to temperature changes above a *T*_*c*_-determined threshold referred to as *T\**_*AFD*_ (Clark et al., 2006; Kimura et al., 2004). Confirming and extending previous observations using different temperature stimulus paradigms (Biron et al., 2006; Matsuyama and Mori, 2020; Ramot et al., 2008a; Tsukada et al., 2016), we found that responses in AFD were largely indifferent to feeding state, although we noted a small but statistically significant decrease in *T\**_*AFD*_ upon starvation (Figure 1E).

Since internal state can modulate downstream circuit components without altering responses in the primary sensory neurons themselves (Datta et al., 2008; Haga et al., 2010; Inagaki et al., 2012; Marella et al., 2012; Rengarajan et al., 2019; Root et al., 2011), we next tested whether temperature responses in AIY, the primary postsynaptic partners of AFD, are altered upon starvation. In a recent study, food deprivation has been suggested to alter the phase relationship between AFD and AIY temperature responses in a restricted temperature range and may mediate plasticity in positive thermotaxis (Matsuyama and Mori, 2020). In contrast to the robust and deterministic temperature responses observed in AFD, we observed stochastic calcium transients in AIY neurons expressing GCaMP6s in immobilized animals in response to the shallow temperature ramp (Figure 1F). We observed no significant differences in total response duration per neuron or the average duration of individual response bouts in AIY between fed and starved animals (Figure 1G, Figure S1C). Moreover, the proportion of AIY neurons responding to the rising temperature stimulus in the temperature range of the behavioral assay (23°C-28°C) was similar under fed and starved conditions, although we noted that responses were initiated at a lower temperature in a subset of AIY neurons upon starvation (Figure 1F). We infer that neither AFD thermosensory responses, nor AFD synaptic output as measured by responses in AIY, are sufficiently modulated by starvation to disrupt thermotaxis, and that alternate pathways must incorporate internal state information elsewhere in the thermotaxis circuit.

### The AWC olfactory neurons integrate feeding state information into the thermotaxis circuit

We and others previously showed that in addition to AFD, the AWC sensory neurons respond to temperature, albeit in a manner distinct from responses in AFD (Biron et al., 2008; Kuhara et al., 2008). However, the contribution of AWC to thermotaxis behaviors is relatively minor under standard assay conditions (Beverly et al., 2011; Biron et al., 2008; Kuhara et al., 2008; Luo et al., 2014a), raising the question of the role of this neuron type in these behaviors. AWC responds robustly to food-related volatile odors (Bargmann et al., 1993; Chalasani et al., 2007), and AWC olfactory responses and AWC-driven behaviors are modulated by the presence or absence of bacterial food (Chalasani et al., 2010; Cho et al., 2016; Colbert and Bargmann, 1995; Lin et al., 2010; Neal et al., 2015; Torayama et al., 2007). We hypothesized that AWC may integrate internal feeding state information into the thermotaxis circuit to modulate negative thermotaxis.

To test this notion, we acutely silenced AWC in adult animals via cell-specific expression of the *Drosophila* histamine-gated chloride channel (HisCl1) and exposure to exogenous histamine (Pokala et al., 2014). Exposing animals expressing HisCl1 in AWC to histamine during the 3h starvation period, but not during the thermotaxis assay, had no effect on the expected negative thermotaxis bias in fed and starved animals (Figure 2A). However, silencing AWC during the assay alone was sufficient to restore negative thermotaxis bias in starved animals (Figure 2A). Moreover, HisCl1-mediated silencing of AWC restored the ability of starved animals to bias the direction of runs following turns towards cooler temperatures (Figure S2A). We confirmed that AWC activity is inhibited by these manipulations by examining AWC-driven olfactory behaviors of animals expressing AWCp::HisCl1. In the presence of histamine, these animals were no longer attracted towards a point source of the AWC-sensed volatile chemical isoamyl alcohol, although they continued to respond to a point source of the odorant diacetyl sensed by the AWA olfactory neuron type (Figure S2B) (Bargmann et al., 1993). Similar HisCl1-mediated silencing of the ASI bacteria-sensing neurons (Gallagher et al., 2013) had no effect on the expected negative thermotaxis behaviors (Figure S2C). As an independent verification, we silenced AWC via cell-specific expression of the light-gated anion channelrhodopsin GtACR2 (Govorunova et al., 2015; Lopez-Cruz et al., 2019). Optogenetic silencing of AWC during the assay was again sufficient to restore negative thermotaxis bias in starved animals (Figure 2B). These results indicate that acute silencing of AWC in adult animals is sufficient to abolish starvation-dependent thermotaxis behavioral plasticity. Moreover, we infer that AWC activity is necessary during the execution of thermotaxis behavior to disrupt negative thermotaxis.

**Figure 2.**
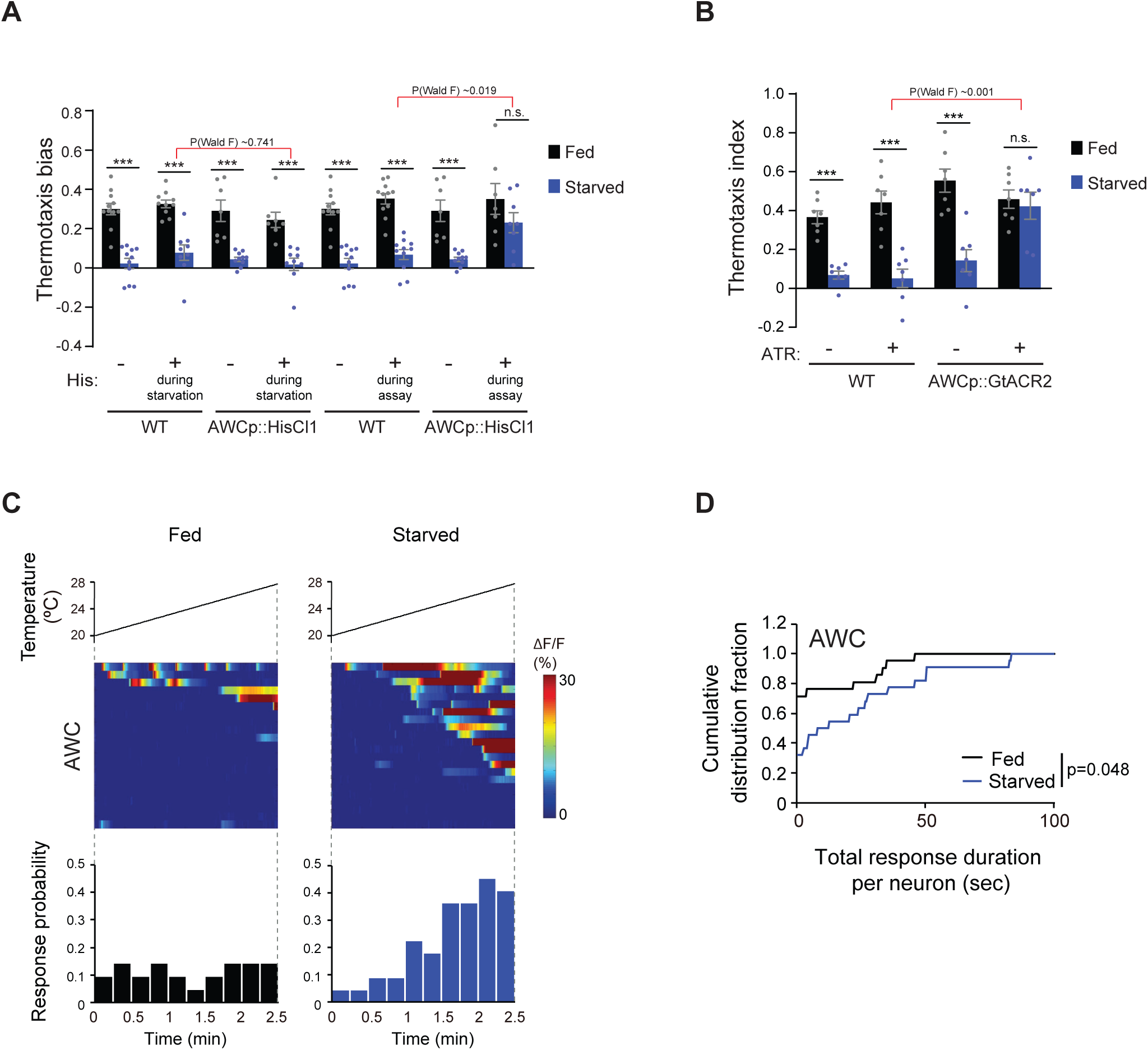
AWC activity is necessary for starvation-dependent suppression of negative thermotaxis. **A)** Mean thermotaxis bias of fed and starved wild-type and transgenic animals expressing HisCl1 in AWC under the *odr-1* promoter in the presence or absence of 10 mM histamine (His). Histamine was present during starvation but not on the assay plate, or only on the assay plate as indicated. Each dot represents the thermotaxis bias of a biologically independent assay comprised of 15 animals. Errors are SEM. *** indicates different from fed at each condition at p<0.001 (Student’s t-test). n.s. – not significant. P-values in red indicate Wald F-statistic from linear regression analysis for the effect of the indicated genotype on the magnitude of the feeding state effect in the indicated conditions. Wild-type data were interleaved with experimental data in Figures 3F-G, and Figure S2C, and are repeated. **B)** Mean thermotaxis index of wild-type and transgenic animals expressing GtACR2 in AWC under the *ceh-36(prom3)* promoter (gift from Steve Flavell). Thermotaxis index was calculated as [(number of animals at 23-24°C on gradient)-(number of animals at 27-28°C on gradient)]/(total number of animals). Animals were grown overnight and assayed with or without 50 □M all-trans retinal (ATR) in the plates as indicated. Assays were performed in the presence of blue light (see Methods). Each dot represents the thermotaxis index of a biologically independent assay comprised of 15 animals. Errors are SEM. *** indicates different from fed at each condition at p<0.001 (Student’s t-test). n.s. – not significant. P-values in red indicate Wald F-statistic from linear regression analysis for the effect of the indicated genotype on the magnitude of the feeding state effect. Wild-type data were interleaved with experimental data in Figure 3H, and are repeated. **C)** Intracellular calcium dynamics in AWC expressing GCaMP3 in response to a linear rising temperature stimulus (black lines) at 0.05°C/sec. Each row in the heatmaps displays responses from a single neuron ordered by the time of the first response; n = 21 (fed) and 22 (starved). (Bottom) Each bar in the histograms represents the percentage of neurons responding during 15s bins. **D)** Cumulative distribution fraction plots of total duration of calcium responses per AWC neuron calculated from data shown in **C**. Distributions were compared using the Kolmogorov-Smirnov test. Also see Figure S2.

We next asked whether temperature responses in AWC are modulated as a function of feeding state. Since the thermal gradient in our behavioral assay for negative thermotaxis ranges from 23°C-28°C (see Methods), we restricted our analysis to this temperature regime. AWC neurons in fed animals grown overnight at 20°C exhibited relatively infrequent and stochastic calcium transients in response to a shallow rising temperature stimulus (Figure 2C) (Biron et al., 2008). We found that the total response duration per neuron as well as the average duration of individual events was significantly increased upon starvation for 3h (Figure 2D, Figure S2D). Moreover, a greater proportion of AWC neurons responded as the temperature increased in starved animals (Figure 2C). The bilateral pair of AWC neurons is functionally asymmetric (AWC^ON^ and AWC^OFF^ neurons) and expresses distinct sets of signaling genes (Troemel et al., 1999). We did not detect obvious asymmetry in the temperature responses of these neurons in starved animals (Figure S2E); these neurons are thus considered together in all subsequent experiments. Together, these results indicate that starvation increases temperature responses in AWC, and that this increased activity is necessary to disrupt negative thermotaxis.

### AWC-mediated inhibition of AIA interneuron temperature responses is necessary and sufficient to mediate starvation-dependent thermotaxis plasticity

How might enhanced temperature responses in AWC in starved animals disrupt negative thermotaxis? Increasing AWC activity either via genetic or optogenetic means has previously been shown to promote reversals and turns via inhibition and activation of the postsynaptic primary layer interneurons AIY and AIA, and AIB, respectively (Figure 3A) (Albrecht and Bargmann, 2011; Chalasani et al., 2007; Chalasani et al., 2010; Gordus et al., 2015; Gray et al., 2005; White et al., 1986). Consequently, AIY and AIA inhibit, and AIB promotes, reversals and/or turns (Figure 3A) (Chalasani et al., 2007; Gordus et al., 2015; Gray et al., 2005; Kocabas et al., 2012; Li et al., 2014; Lopez-Cruz et al., 2019; Tsalik and Hobert, 2003; Wakabayashi et al., 2004). Enhanced AWC activity in starved animals is consistent with our observation that animals exhibit increased reversals and turns at the starting temperature upon food deprivation (Figure 1B), thereby failing to correctly navigate the spatial thermal gradient. We asked whether AWC acts via one or more of these first layer interneurons to regulate negative thermotaxis.

**Figure 3.**
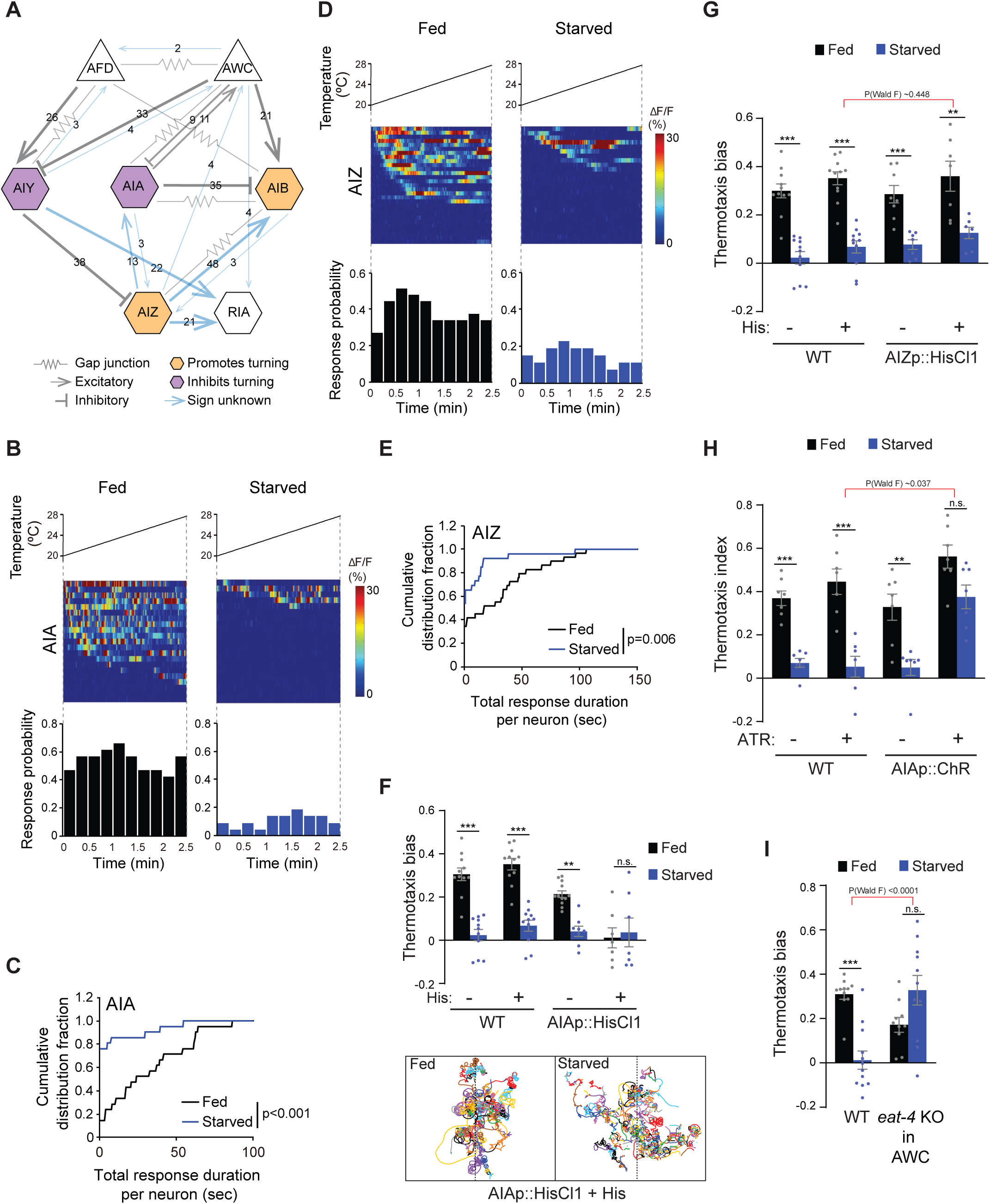
Inhibition of AIA temperature responses is necessary and sufficient for starvation-dependent suppression of negative thermotaxis. **A)** Schematic of chemical and electrical connectivity of indicated sensory neurons and interneurons. Connections whose signs have not been experimentally validated are indicated in blue. Numbers of synapses (>2) observed via serial section electron microscopy are indicated. Weights of connecting lines are scaled to synaptic strength. Color codes indicate neurons whose activity is associated with promotion or inhibition of reversals and turns. Adapted from (Cook et al., 2019; White et al., 1986) (www.wormwiring.org). **B, D)** Intracellular calcium dynamics in AIA **(B)** and AIZ **(D)** expressing GCaMP5A (AIA) or GCaMP6s (AIZ) in response to a linear rising temperature stimulus (black lines) at 0.05°C/sec. Each row in the heatmaps displays responses from a single neuron ordered by the time of the first response; n = 21 (AIA, each fed and starved), 29 (AIZ, fed) and 26 (AIZ, starved). (Bottom) Each bar in the histograms represents the percentage of neurons responding during 15s bins. **C, E)** Cumulative distribution fraction plots of total duration of calcium responses per AIA **(C)** and AIZ **(E)** neuron calculated from data shown in **B** and **D**, respectively. Distributions were compared using the Kolmogorov-Smirnov test. **F, G)** Mean thermotaxis bias of fed and starved wild-type and transgenic animals expressing HisCl1 in AIA **(F)**, and AIZ **(G)** in the presence or absence of 10 mM histamine on the assay plate (see Table S1 for genotypes). Each dot represents the thermotaxis bias of a biologically independent assay comprised of 15 animals. Errors are SEM. *** and ** indicate different from fed at each condition at p<0.001 and p<0.01, respectively (Student’s t-test). n.s. – not significant. Wild-type data were interleaved with experimental data in Figures 2A, and Figure S2C, and are repeated. P-values in red indicate Wald F-statistic from linear regression analysis for the effect of the indicated genotype on the magnitude of the feeding state effect. Traces in F show trajectories of fed and starved animals from a representative 35 min assay of 15 animals expressing HisCl1 in AIA on histamine-containing plates; dashed lines indicate the starting temperature of 25.5°C on the thermal gradient. Individual worm trajectories are color-coded. Since trajectories are terminated by omega turns or collisions, trajectories of the same color may not represent the movement of a single animal throughout the assay. **H)** Mean thermotaxis index of wild-type and transgenic animals expressing ChR in AIA under the *ins-1(s)* promoter. Animals were grown overnight and assayed with or without 50 □M all-trans retinal (ATR) in the plates as indicated. Assays were performed in the presence of blue light. Each dot represents the thermotaxis index of a biologically independent assay comprised of 15 animals. Errors are SEM. *** and ** indicate different from fed at each condition at p<0.001 and p<0.01, respectively (Student’s t-test). n.s. – not significant. P-values in red indicate Wald F-statistic from linear regression analysis for the effect of the indicated genotype on the magnitude of the feeding state effect. Wild-type data were interleaved with experimental data in Figure 2B, and are repeated. **I)** Mean thermotaxis bias of animals of the indicated genotypes. *eat-4* was knocked out in AWC via FLP-FRT-mediated recombination (Lopez-Cruz et al., 2019). Each dot represents the thermotaxis bias of a biologically independent assay comprised of 15 animals. Errors are SEM. *** indicates different from fed p<0.001 (Student’s t-test). n.s. – not significant. P-values in red indicate Wald F-statistic from linear regression analysis for the effect of the indicated genotype on the magnitude of the feeding state effect. Wild-type control data were interleaved with experimental data in Figures S3E, Figure S4C, and S4F, and are repeated. Also see Figure S3.

Since AIY temperature responses are largely indifferent to feeding state (Figure 1F), we examined temperature responses in AIA and AIB in immobilized fed and starved animals. Similar to AIY, both AIA and AIB exhibited stochastic calcium transients in response to a shallow rising temperature stimulus in the *T>T*_*c*_ regime (Figure 3B, Figure S3A). AIB temperature responses were similar in fed and starved animals (Figure S3A-B). AIB activity is tightly coupled to the motor state of animals (Gordus et al., 2015; Kato et al., 2015). The observation that AIB activity is unaltered in starved animals despite changes in locomotory output raises the possibility that under these conditions, AIB may be partly disassociated from network activity state. However, temperature responses in AIA were markedly altered upon starvation (Figure 3B-C, Figure S3C). AIA appeared to be tonically active and exhibited frequent calcium transients in fed animals in response to the rising temperature stimulus; the proportion of responding animals, the total response duration per neuron as well as the average duration of individual response bouts were significantly suppressed upon prolonged starvation (Figure 3B-C, Figure S3C). Temperature responses in the AIZ interneurons have also previously been reported to be modulated by feeding state (Kodama et al., 2006). Although not directly postsynaptic to AWC (Figure 3A), AIZ is a major postsynaptic partner of AIY, receives inputs from both AIA and AIB and multiple sensory neuron types, and is presynaptic to RIA (Cook et al., 2019; White et al., 1986) (Figure 3A). Under our imaging conditions, temperature responses in AIZ, like those in AIA, exhibited tonic stochastic activity in fed animals subjected to a rising temperature stimulus, and these responses were decreased upon starvation (Figure 3D-E, Figure S3D).

We next asked whether suppression of AIA and/or AIZ activity in fed animals is sufficient to disrupt negative thermotaxis. To inhibit AIA, we expressed HisCl1 under the *gcy-28d* promoter which drives expression strongly in AIA and less consistently in AVF and a subset of additional neurons (the ‘AIA circuit’) (Cho et al., 2016). Fed and starved animals expressing HisCl1 in the AIA circuit continued to exhibit the expected negative thermotaxis bias in the absence of histamine (Figure 3F). However, acute inhibition of the AIA circuit via addition of histamine to the assay plate resulted in animals exhibiting extensive reversals and turns at the starting temperature regardless of feeding state, and consequent inability of these animals to navigate the gradient (Figure 3F). We observed similar effects on animal locomotion upon inhibition of AIA via expression of an activated UNC-103 potassium channel [*unc-103(gf)*] in AIA (Cho et al., 2016; Lin et al., 2010; Reiner et al., 2006) (Figure S3E). However, although acute inhibition of AIZ via expression of HisCl1 and addition of histamine decreased spontaneous reversals on an isothermal plate as reported previously (Gray et al., 2005; Li et al., 2014; Tsalik and Hobert, 2003) (Figure S3F), AIZ inhibition had no effect on the expected thermotaxis behavior of either fed or starved animals (Figure 3G). We conclude that while starvation inhibits temperature responses in both AIA and AIZ, complete suppression of these responses in AIA but not AIZ disrupts negative thermotaxis regardless of feeding state.

We next asked whether acute activation of AIA is sufficient to restore negative thermotaxis in starved animals. We optogenetically activated AIA via expression of the light-activated ion channel Chrimson under the *ins-1(s)* promoter (Dobosiewicz et al., 2019; Klapoetke et al., 2014) as animals performed thermotaxis on a spatial thermal gradient. Activation of AIA was sufficient to restore the ability of starved animals to perform negative thermotaxis (Figure 3H). AWC inhibits AIA via glutamatergic signaling (Chalasani et al., 2010). If AWC-mediated glutamatergic transmission inhibits temperature responses in AIA in starved animals, we would predict that blocking glutamatergic signaling from AWC would also be sufficient to restore negative thermotaxis upon starvation. Indeed, we found that knocking out the glutamate transporter *eat-4* cell-specifically in AWC (Lopez-Cruz et al., 2019) resulted in robust negative thermotaxis behavior by starved animals (Figure 3I). These experiments suggest that upon prolonged starvation, increased AWC temperature responses inhibit AIA via glutamatergic signaling to disrupt negative thermotaxis via warming-uncorrelated regulation of reversals and turns. These observations also provide a mechanistic explanation for the previously reported decorrelation between AFD activity and turns in starved as compared to fed animals navigating a thermal gradient (Tsukada et al., 2016). Moreover, these observations indicate that suppression or activation of AIA in fed or starved animals is sufficient to permit or inhibit negative thermotaxis, respectively.

### INS-1 signaling from the intestine regulates thermotaxis behavioral plasticity in response to starvation

INS-1 ILP signaling has previously been implicated in feeding state-dependent modulation of negative thermotaxis (Kodama et al., 2006), although the source and target of this signaling are unclear. *ins-1* has previously been shown to be expressed in multiple neuron types, including AIA as well as in the intestine, as assessed via GFP reporter expression under *ins-1* regulatory sequences (Kodama et al., 2006; Pierce et al., 2001; Tomioka et al., 2006). We investigated the required source of INS-1 production for the modulation of negative thermotaxis in starved animals. To address this issue, we knocked out *ins-1* cell-specifically using Cre-Lox-mediated recombination. We generated strains carrying *loxP* sites flanking the endogenous *ins-1* locus as well as extrachromosomal arrays driving expression of Cre tagged with GFP under the *ins-1* endogenous or cell-specific promoters (Figure S4A). We then selected animals expressing Cre::GFP in the cells of interest and examined their thermotaxis behaviors under fed and starved conditions. Knocking out *ins-1* in all or a majority of *ins-1*-expressing cells, but not in AIA or AIZ/AIB, was sufficient to restore negative thermotaxis in starved animals (Figure 4A). We found that knocking out *ins-1* specifically in the intestine phenocopied the negative thermotaxis behavior of *ins-1* null mutants (Figure 4A, 4D) (Kodama et al., 2006), indicating that *ins-1* production from the intestine is necessary for starvation-mediated plasticity in negative thermotaxis.

**Figure 4.**
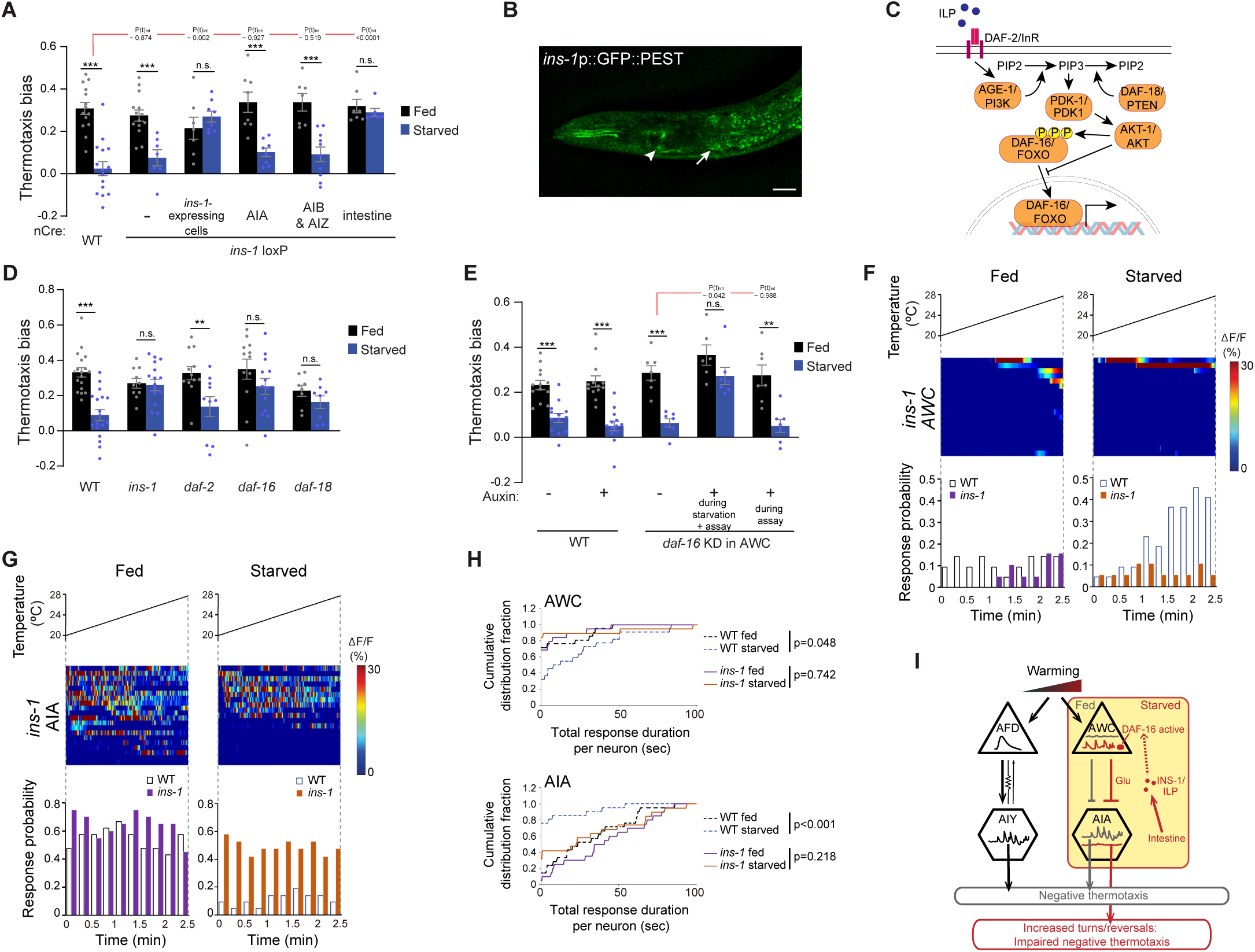
INS-1 insulin signaling from the gut modulates AWC and AIA temperature responses to disrupt negative thermotaxis upon starvation. **A, D, E)** Mean thermotaxis bias of animals of the indicated genotypes. *ins-1* was knocked out cell-specifically via Cre-Lox-mediated recombination using *ins-1* alleles flanked with *loxP* sequences, and cell-specific expression of nCre **(A)** (Figure S4A, Table S1). Alleles used in **(D)** were *ins-1(nr2091), daf-2(e1368), daf-16(m26)*, and *daf-18(ok480)*. DAF-16 was depleted in AWC via auxin-induced degradation of a degron-tagged *daf-16* allele and AWC-specific expression of TIR1 under the *ceh-36prom2_del1*ASE promoter **(E)** (Figure S4D). Auxin was added during starvation and to the assay plate, or to the assay plate alone, as indicated in **E**. Each dot represents the thermotaxis bias of a biologically independent assay comprised of 15 animals. Errors are SEM. *** and ** indicate different from fed p<0.001 and p<0.01, respectively (Student’s t-test). n.s. – not significant. P-values in red indicate t-statistic from posthoc effect size comparisons (Dunnett’s test) between the indicated genotypes **(A)** or the conditions **(E)**, respectively, on the magnitude of the feeding state effect. Wild-type data in **E** were interleaved with experimental data in Figure S4E, and are repeated. **B)** Representative image of the expression pattern of an *ins-1*p::GFP::PEST reporter in a well-fed adult hermaphrodite. Expression in the gut and in AIA are indicated by an arrow and arrowhead, respectively. Anterior is at left. Scale bar: 20 □m. **C)** Cartoon of the canonical insulin signaling pathway. **F, G)** Intracellular calcium dynamics in AWC **(F)** and AIA **(G)** neurons in *ins-1* mutants expressing GCaMP3 (AWC) or GCaMP5A (AIA) in response to a linear rising temperature stimulus (black lines) at 0.05°C/sec. Each row in the heatmaps displays responses from a single neuron ordered by the time of the first response; n = 19 (AWC, each fed and starved), 20 (AIA, fed) and 19 (AIA, starved). (Bottom) Each bar in the histograms represents the percentage of neurons from animals of the indicated genotypes responding during 15s bins. Open bars indicate data from wild-type neurons repeated from Figure 2C (AWC) and Figure 3B (AIA). **H)** Cumulative distribution fraction plots of total duration of calcium responses per AWC (top) and AIA (bottom) neuron calculated from data shown in **F** and **G**, respectively. Dashed lines indicate wild-type data repeated from Figure 2D (AWC) and Figure 3C (AIA). Distributions were compared using the Kolmogorov-Smirnov test. **I)** Working model of AWC- and AIA-mediated disruption of negative thermotaxis upon starvation. AFD and AIY temperature responses are indifferent to feeding state. In fed animals, low and high temperature responses in AWC and AIA, respectively is permissive for AFD-mediated negative thermotaxis. Upon prolonged starvation, INS-1 signaling from the gut acts directly or indirectly on AWC via DAF-16 to increase temperature responses. Glutamatergic signaling from AWC inhibits temperature responses in AIA resulting in temperature change-uncorrelated reversals and turns and disruption of AFD-mediated negative thermotaxis. Also see Figure S4.

We asked whether prolonged starvation alters *ins-1* expression in the intestine by examining expression of destabilized GFP (GFP::PEST) reporter driven under the *ins-1* promoter. As reported previously, we observed expression of this reporter in head neurons including in AIA as well as in the intestine (Figure 4B). However, levels of *ins-1*p::GFP::PEST expression in the gut were not altered upon food deprivation for 3h (Figure S4B), suggesting that mechanisms other than changes in intestinal *ins-1* expression account for the effects of prolonged food deprivation on negative thermotaxis.

### INS-1 targets AWC to alter temperature responses as function of food deprivation

ILPs act as agonists or antagonists of the DAF-2 insulin receptor to regulate the phosphorylation state and subcellular localization of the DAF-16 FOXO transcription factor (Figure 4C) (Murphy and Hu, 2013; Tissenbaum, 2018). Unlike *ins-1* mutants but similar to wild-type animals, animals mutant for *daf-2*, or the *age-1* and *akt-1* kinases in the insulin signaling pathway failed to perform negative thermotaxis upon starvation (Figure 4D, Figure S4C). We could not examine the effects of mutations in the *pdk-1* kinase on thermotaxis since these mutants were athermotactic regardless of feeding state (Figure S4C). Loss of function of both the *daf-18* PTEN phosphatase and *daf-16* FOXO restored the ability of starved animals to perform negative thermotaxis (Figure 4D). We interpret these results to indicate that while the presence or absence of DAF-2-mediated signaling does not influence negative thermotaxis in fed animals, INS-1 directly or indirectly antagonizes DAF-2 signaling and disrupts negative thermotaxis via promotion of DAF-16 activity.

We asked whether intestinally-produced INS-1 targets AWC to alter thermotaxis behavior in starved animals by cell-specifically depleting DAF-16 protein in AWC using the auxin-inducible degron system (Zhang et al., 2015). We expressed the TIR1 F-box protein specifically in AWC in animals in which the *daf-16* locus is genome-engineered with auxin-inducible degron sequences (Figure S4D) (Aghayeva et al., 2020). We found that addition of auxin during the 3h starvation period and the assay but not during the assay alone resulted in significant rescue of negative thermotaxis behavior (Figure 4E). Although it is possible that DAF-16 is not sufficiently degraded upon addition of auxin only during the assay period, 4mM auxin was previously shown to reduce degron-tagged protein abundance to <5% within 30 mins (Zhang et al., 2015). We did not observe any effects on expected negative thermotaxis behavior upon auxin-mediated DAF-16 degradation specifically in ASI (Figure S4E). Moreover, we observed no changes in the expected negative thermotaxis behaviors of animals mutant for additional genes previously implicated in feeding state-dependent modulation of AWC-driven olfactory behaviors (Figure S4F) (Chalasani et al., 2010; Neal et al., 2015; Tsunozaki et al., 2008). We infer that INS-1 targets DAF-16 in AWC to modulate negative thermotaxis bias as a function of feeding state, and that DAF-16 function is necessary during prolonged starvation for this modulation.

We next asked whether starvation-regulated plasticity in temperature responses in AWC and AIA was affected in *ins-1* mutants. Indeed, we found that in contrast to the increased activity in AWC observed in starved wild-type animals, temperature responses is this neuron type were no longer modulated by feeding state in *ins-1* mutants (Figure 4F, 4H, Figure S4G). Specifically, AWC neurons in both fed and starved *ins-1* mutants exhibited infrequent responses similar to the response profiles of fed wild-type animals (Figure 4F, 4H, Figure S4G). Similarly, the proportion of responding animals, the total response duration per neuron, as well as the average duration of individual events in AIA in fed and starved *ins-1* mutants resembled the activity profiles of AIA neurons in fed wild-type animals (Figure 4G-H, Figure S4H). We conclude that INS-1 signaling from the intestine in starved animals alters AWC and AIA temperature responses to disrupt negative thermotaxis.

## Discussion

We show here that prolonged starvation disrupts negative thermotaxis by altering the activity of the modulatory AWC-AIA, but not that of the core thermosensory AFD-AIY, circuit. In fed animals, warming fails to evoke responses in AWC, and elicits temperature responses in AIA. Under these conditions, AIA-mediated inhibition of reversals and turns enables AFD-driven regulation of klinokinesis and turning bias as a function of temperature changes and the animal’s *T*_*c*_ (Figure 4I). However, during prolonged food deprivation, INS-1 signaling from the gut acts via DAF-16 in AWC neurons to alter their response properties. The altered state of AWC in starved animals results in enhanced and reduced temperature responses in AWC and AIA, respectively, and consequent disruption of AFD-driven thermotactic navigation (Figure 4I). Our results indicate that the activity state of AWC-AIA is permissive for AFD-dependent negative thermotaxis in fed animals, and that food deprivation modulates AWC-AIA via gut-to-brain signaling to disrupt this behavior. Starvation has previously been shown to enhance odorant responses in AWC (Neal et al., 2015). We speculate that starvation-dependent changes in sensory responses in AWC allow *C. elegans* to prioritize responses to bacterial food-related odorants over optimal thermoregulation.

We find that INS-1 signaling from the gut is necessary to modulate AWC responses upon prolonged starvation. In contrast, *ins-1* expression in AIA has been shown to be sufficient for modulation of chemosensory responses in AWC as well as in the ASER sensory neurons upon pairing starvation with AWC- or ASE-sensed chemical cues (Chalasani et al., 2010; Cho et al., 2016; Lin et al., 2010; Tomioka et al., 2006). Similarly, expression of *ins-1* from multiple neuron types rescues the *ins-1* thermotaxis behavioral phenotype in starved animals (Kodama et al., 2006). These observations suggest that while *ins-1* expression from multiple neuronal sources including AIA is sufficient, expression and/or production from specific cells and tissues, potentially at different levels, and in response to changing internal states over time, may be necessary for neuromodulation in AWC. The expression of multiple ILP genes as well as neuropeptides is regulated by diverse environmental and internal conditions in different cell types in *C. elegans* (Cornils et al., 2011; Fernandes de Abreu et al., 2014; Kim and Li, 2004; Pierce et al., 2001; Ritter et al., 2013). Given the possibility of ILP-to-ILP signaling (Chen et al., 2013; Fernandes de Abreu et al., 2014), intestinal INS-1 may target AWC indirectly via other ILPs expressed from neuronal or non-neuronal tissues. It is likely that regulated transcription as well as release of ILPs and other peptides from defined cell and tissue types are critical for modulation of specific neuron types as a function of temporally varying internal states.

As we reported previously (Beverly et al., 2011; Biron et al., 2008), AWC responses to a shallow rising temperature stimulus are stochastic, although their duration is stimulus-regulated. However, AWC responses to a steeply rising temperature stimulus are time-locked (Kuhara et al., 2008). Similarly, while the ASER salt-sensing neurons exhibit stochastic activity in response to a shallow linear salt gradient, they exhibit time-locked responses when subjected to large step changes in salt concentrations (Luo et al., 2014b). Large stimulus changes may saturate intracellular calcium levels and mask physiologically relevant underlying neuronal dynamics. While AFD exhibits extraordinary thermosensitivity, temperature responses in AWC appear to be far less sensitive (Ramot et al., 2008a). We suggest that a shallow thermal ramp may not be sufficient to elicit sustained temperature responses in AWC, resulting in the observed stochastic activity pattern. Upon prolonged starvation, DAF-16-regulated expression changes in thermosensory signaling molecules may lead to the observed increased frequency and duration of temperature response in AWC. Molecules such as the CMK-1 calcium/calmodulin-dependent protein kinase I and the EGL-4 cGMP-dependent protein kinase have previously been implicated in regulating AWC olfactory responses as a function of starvation in distinct contexts (Cho et al., 2016; Lee et al., 2010; Neal et al., 2015), suggesting that distinct pathways may translate internal state information into changes in AWC responses under different conditions. An important next step will be to identify targets of these pathways in AWC in animals subjected to defined experiences.

Interestingly, although temperature responses in AIZ resemble those in AIA in fed and starved animals, inhibition of AIZ has no effect on negative thermotaxis under either fed or starved conditions. The circuit underlying thermotaxis behaviors in *C. elegans* is remarkably degenerate, and alternate neuronal pathways can compensate for the absence of the core AFD and AIY components in distinct contexts for different aspects of thermotaxis behaviors (Beverly et al., 2011; Matsuyama and Mori, 2020). AIZ may be a component of a degenerate circuit that modulates negative thermotaxis under specific environmental or genetic conditions. Although this hypothesis remains to be verified, the presence of degenerate circuits driving specific behaviors in a context-dependent manner is a conserved feature of nervous systems, and contributes to both robustness and flexibility in behavioral outputs (Cropper et al., 2016; Edelman and Gally, 2001; Prinz et al., 2004; Saideman et al., 2007; Trojanowski et al., 2014; Wang et al., 2019).

Acute inhibition of AWC and activation of AIA is sufficient to restore negative thermotaxis in starved animals, indicating that the functions of the core thermotaxis circuit are maintained, but are masked by the activity state of the AWC-AIA circuit. What might be the advantage of driving behavioral plasticity via this gating mechanism as compared to direct regulation of the core circuit? We suggest that incorporating distinct modulatory pathways as a function of different experiences and conditions allows for a greater degree of behavioral flexibility, as compared to direct modulation of the core circuit itself, particularly in small circuits. Food deprivation for different periods of time has been shown to result in distinct behavioral changes that may be physiologically relevant in terms of driving specific food seeking behaviors (Churgin et al., 2017; Farhadian et al., 2012; Ghosh et al., 2016; Gray et al., 2005; Inagaki et al., 2014; Lee and Park, 2004; Lopez-Cruz et al., 2019; Neal et al., 2015; Root et al., 2011; Tsalik and Hobert, 2003). These changes may be driven via temporally regulated recruitment of different neuronal pathways to functionally reconfigure the core sensorimotor circuit and reprioritize behaviors (Churgin et al., 2017; Ghosh et al., 2016; Gray et al., 2005; Inagaki et al., 2012; Saeki et al., 2001). Although modulatory pathways may target different nodes in the underlying circuits, sensory neurons and first layer interneurons are the primary targets in *C. elegans* presumably due to the relatively shallow network architecture of the nematode nervous system.

Functional reconfiguration of circuits via neuromodulation expands the repertoire of circuit outputs (Bargmann, 2012; Griffith, 2013; Kim et al., 2017; Marder, 2012). The mechanisms by which neuromodulators effect plasticity in circuit and behavioral output are diverse. Neuromodulation can alter sensory or synaptic gain in specific pathways, enable integration of neurons and pathways into defined functional circuits, selectively activate or inhibit a subset of available hardwired synaptic connections, and broadly regulate circuit state to affect excitability [eg. (Baldridge et al., 1998; Cho et al., 2016; Cohn et al., 2015; Ha et al., 2010; Hill et al., 2015; Jing et al., 2007; Komuniecki et al., 2014; Macosko et al., 2009; Marder and Hooper, 1985)]. Identification of the pathway by which internal state regulates thermotaxis behavioral plasticity in *C. elegans* adds to our understanding of the mechanistic richness of neuromodulator functions, and suggests that related principles may operate across diverse sensory circuits.

## Material and Methods

### *C. elegans* strains

The wild-type strain used was *C. elegans* variety Bristol strain N2 grown on *E. coli* OP50. Transgenic animals were generated using experimental plasmids at 2-50 ng/μl and the *unc-122*p::*gfp* or *unc-122*p::*dsRed* coinjection markers at 30 ng/μl unless noted otherwise. Expression patterns and behavioral phenotypes were confirmed in initial experiments using multiple independent transgenic lines, and typically, a single line was selected for additional analysis. The presence of specific mutations and genome edits were confirmed by sequencing. A complete list of all strains used in this work is provided in Table S1.

To express *TiR1::mTurquoise2* in individual neurons, 15 ng/μl of *TiR1* sequences under cell-specific promoters were injected together with 100 ng/μl of linearized N2 genomic DNA, and 30 ng/μl of co-injection marker. To stably integrate *ins-1*p::*gfp::PEST* sequences into the genome, young transgenic adults carrying extrachromosomal arrays were irradiated with 300 μJ UV light (UVP Imaging System). Animals homozygous for the integrated array were backcrossed 3 times prior to use.

### Molecular biology

Promoter sequences and cDNAs were amplified from plasmids or a *C. elegans* cDNA library generated from a population of mixed stage animals. Promoters used in this work were (upstream of ATG): *odr-1* (AWC: 1.0 kb), *srg-47* (ASI: 650 bp), *ins-1* (4.2 kb), *srsx-3* (AWC^OFF^: 1.3 kb), *gcy-28d* (AIA circuit: 2.8 kb), *odr-2b(3a)* (AIZ and others: 448 bp), *ser-2(2)* (AIZ and others), and *ifb-2* (intestine: 3.0 kb). GFP-PEST sequences were amplified from an *nlp-36*p::GFP::PEST encoding plasmid (van der Linden et al., 2010). The plasmid containing *ceh-36prom2_del1*ASEp::*Tir1::mTurquiose2* was a gift from the Hobert lab. The *odr-2b(3a)* promoter-containing plasmid was a gift from Shawn Xu. Plasmids were generated using Gibson assembly (New England BioLabs) or In-Fusion cloning (Takara Bio) unless noted otherwise.

To generate WY016, *FRT::STOP::FRT* sequences were amplified from sequences in the *ser-2(2)*p::*FRT::YFP* plasmid (gift from Shawn Xu), and inserted into a vector containing *GCaMP6s* sequences (Wu et al., 2019) by Gibson assembly (New England BioLabs). *ser-2(2)*p (4.7 kb) sequences were amplified from plasmids derived from *ser-2(2)*p::*FRT::YFP*,andclonedupstreamof *FRT::STOP::FRT::GCaMP6s* to generate WY016 using Gateway (Thermo Fisher Scientific). To generate PSAB1206, *HisCl1::SL2::mCherry* sequences from PSAB1204 were inserted downstream of *ser-2(2)*p::*FRT::STOP::FRT*. All plasmids generated and used in this work are listed in Table S2.

### Generation of *ins-1(oy158)*

To generate *ins-1(oy158), loxP* sequences were sequentially inserted 50 bp and 534 bp upstream and downstream, respectively, of the *ins-1* locus via gene editing (Figure S4A). Donor oligonucleotides (IDT: Integrated DNA Technologies) containing 35 bp homology arms (Dokshin et al., 2018) were injected (110 ng/μl) together with crRNA (20ng/μl; IDT), tracrRNA (20ng/μl, IDT), Cas9 protein (25 ng/μl; IDT), and *unc-122*p::*dsRed* (30ng/μl) as the co-injection marker. F1 animals expressing the injection marker were isolated, and genome editing was confirmed by amplification and sequencing. F2 progeny were further screened for the presence of the homozygous genome edited sites. To knockout expression of *ins-1* cell-specifically, *nCre::SL2::gfp* sequences were expressed under *ins-1* endogenous (4.2 kb), *ifb-2, gcy-28d*, and *odr-2b(3a)* regulatory sequences and animals expressing nCre in the required cell and tissue types were examined in behavioral assays. We were unable to insert fluorescent reporter sequences together with *loxP* sequences either 5’ or 3’ of the *ins-1* genomic locus in multiple (>7) attempts. Sequences of crRNAs and donor oligonucleotides used were: crRNA(upstream):5’-CTCGGAAATATATATTTATGTTTTAGAGCTATGCT - 3’, crRNA(downstream):5’-TACCATTTATTTCTATAAATGTTTAGAGCTATGCT - 3’ Donoroligo(upstream):5’-CCCGTTGTTGAGAGCGGTGAGGAACTGAAAAATGCATAACTTCGTATAATGTATGCTATACGAAGTTATACCGGTTCTATAAATATATATTTCCGAGTACTAAAAACGAAAACGAA - 3’. Donor oligo(downstream):5’-GTTCAAACTGCGTCACATTTGTGATCAAATGTTGAAAATATTTATAGAAATAAATGGTATAACCGGTAAATAACTTCGTATAGCATACATTATACGAAGTTATTTTTGAATGAATTTTTCAAGGTCGCCGATTTGCCGG - 3’.

### Thermotaxis behavioral assays

*C. elegans* was grown for at least 3 generations at 20°C with ample OP50 bacterial food prior to being examined in behavioral assays. To obtain starved animals, 3h prior to the start of the assay, young adult worms were transferred twice sequentially to unseeded NGM plates and allowed to move freely for a few minutes to remove associated bacteria. Animals were then transferred to fresh unseeded NGM plates and re-cultivated at the appropriate temperature. To expose animals to bacterial odors, unseeded NGM plates containing worms were covered with a seeded agar plate for 3h. To test the necessity of live bacteria, animals were fed with gentamycin-treated OP50 spread on NGM plates for 3h prior to the behavioral assay (200 μg/ml gentamycin was added to concentrated OP50 and incubated for 2h). For re-feeding, animals starved for 3h were transferred to OP50-seeded NGM plates for 1h prior to the assay. Just prior to behavioral assays, 25 animals were transferred to unseeded NGM plates briefly, picked into M9 buffer pre-incubated at the cultivation temperature and subsequently transferred to the thermal gradients. Transfer from the cultivation plate to the thermal gradient was typically accomplished within 5 mins. *Long thermal gradient:* To examine negative and positive thermotaxis on the long gradient (Luo et al., 2014a), worms grown at 20°C were transferred to 15°C or 25°C, respectively, for 4h prior to the assay. The temperature on the aluminum plate (ranging from 18-22°C at 0.2°C/cm) was controlled by a Peltier system [colder side; H-bridge amplifier (Accuthermo FTX700D), PID controller (Accuthermo FTC100D), Peltier (DigiKey] and heater system [warmer side; PID controller(OmegaCNi3244),solid-staterelay(Omega SSRL240DC25), and cartridge heaters (McMaster-Carr 3618K403)]. A 22.5cm square NGM agar pad for the assay was placed directly on the aluminum plate, and the temperature of agar edges and center were confirmed with a two-probe digital thermometer (Fluke Electronics) prior to each assay. Worms in M9 buffer were placed on the 20°C isotherm at the center of the gradient. Animal movement was imaged at 2 fps for 30 min using a Mightex camera (BTE-5050-U). Animal trajectories were detected and analyzed using custom LabView and MATLABscripts(Gershowetal.,2012) (https://github.com/samuellab/MAGATAnalyzer).

#### Short thermal gradient

A ‘short’ thermal gradient was established on an unseeded 10 cm plate containing 25 ml NGM agar placed on an aluminum sheet. The thermal gradient (ranging from 23°C-28°C at 0.5°C/cm) was established and maintained on the aluminum sheet using Peltier thermoelectric temperature controllers (Oven Industries). The temperature of the edges of NGM agar was measured with a two-probe digital thermometer (Fluke Electronics). Worms in M9 buffer cultivated at 20°C were placed at the center of the gradient at 25.5°C. Animal movement was recorded at a rate of 1 Hz using a PixeLink CCD camera controlled by custom written scripts in MATLAB for 35 mins. 30 min videos excluding the first 5 mins were analyzed using WormLab (MBF Bioscience) and custom written scripts in MATLAB as described previously (Beverly et al., 2011; Yu et al., 2014).

#### Histamine-mediated acute inhibition

10 mM histamine-containing plates were generated essentially as described previously (Pokala et al., 2014). 1M histamine dihydrochloride (Sigma-Aldrich H7250) was added to NGM agar at 60°C prior to pouring into Petri plates. Animals were grown on bacteria-seeded or unseeded NGM agar plates containing histamine for 3h prior to the assay as indicated. For histamine-mediated inhibition during the assay, 10 mM histamine was added to the NGM agar plate on which the thermal gradient was established.

#### Optogenetic activation/inhibition

L4 larval animals were fed 50 μM ATR-OP50 overnight. 3h prior to the assay, animals were transferred to seeded or unseeded 50 μM ATR containing NGM plates. Animals were placed on short thermal gradients as described but were also exposed to blue LED light (approximately wavelength 455-470) at 1.9 mW/mm^2^ intensity for 25 mins during the assay. At the completion of the assay, the number of animals at the cold and warm ends of the plate were counted to calculate the thermotaxis index [(number of animals at the temperature range 23-24°C)-(number of animals at 27-28°C)]/(total number of animals in assay).

#### Auxin-induced degradation

400 mM Auxin in EtOH (indole-3-acetic acid, Alfa Aesar) was used to make 4 mM auxin-containing seeded, unseeded or assay plates. Plates were kept wrapped in aluminum foil to prevent light exposure.

#### Analysis of thermotaxis bias and reorientation direction following turns

Identification of worm tracks, direction and duration of worm runs, and run orientation following turns were analyzed with custom written scripts in MATLAB (Beverly et al., 2011; Yu et al., 2014). Each track was defined as a continuous worm trajectory between turns. Track orientation following turns was calculated as the angle of a line connecting the initial and last points of the location of the worm in successive tracks. Thermotaxis bias was calculated as (run duration toward colder side – run duration toward warmer side)/total run duration (Beverly et al., 2011; Chi et al., 2007; Clark et al., 2007). A minimum of seven biologically independent trials with at least 15 animals each were conducted for each experimental condition. Behaviors of mutant and transgenic animals were assessed in parallel with wild-type controls on the same day. Wild-type data were interleaved with data from experimental strains collected over a similar time period and are repeated as indicated in the Figure Legends.

### Olfactory behavioral assays

Chemotaxis assays were performed as previously described (Bargmann et al., 1993). In brief, well-fed animals washed twice with S-Basal and once with water were placed at the center of a 10 cm NGM agar plate with or without 10 mM histamine. 1 μl of isoamyl alcohol (Fisher A393-500) and diacetyl (Sigma B85307) diluted 1:1000 with ethanol were placed at one end with 1 μl of ethanol as the diluent control placed at the other end, together with 1 μl of 1M sodium azide. Animals at either end were counted after 60 mins. The chemotaxis index was defined as [(number of animals at the odorant) – (number of animals at the diluent)/ total number of animals]. For transgenic strains, only transgenic animals as assessed by expression of the coinjection marker were counted. Prior assessment indicated that 98% of animals (n=50) expressing the coinjection marker in the PY12205 strain also expressed *HisCl1::SL2::mCherry* in both AWC neurons.

### Quantification of spontaneous reversal frequency

Young adult animals were transferred onto NGM plates at 20°C containing a thin layer of OP50 bacteria. Reversal frequencies were quantified following 1 min after transfer for 20 mins. Backward movement with two or more head swings were scored manually as reversals (Gray et al., 2005).

### *In vivo* calcium imaging

Calcium imaging experiments were performed essentially as described previously (Takeishi et al., 2016). AWC^ON^ and AWC^OFF^ neurons were identified via expression of *mScarlet* driven under *srsx-3* regulatory sequences. Growth-synchronized L4 larval animals were cultivated overnight with ample OP50 at 20°C. Young adult animals were starved by placing them on unseeded NGM agar plates and re-cultivating at 20°C for 3h prior to imaging.

Individual animals were glued (WormGlu, GluStitch Inc.) to an NGM agar pad on a cover glass, bathed in M9, and mounted under a second cover glass for imaging. For imaging of AFD and AIY, 5-10 fed or starved young adult worms were picked into 10 µM levamisole diluted in M9 on a 5% agarose pad and immobilized between two coverslips. The edges of the cover glass sandwich were sealed with a mixture of paraffin wax (Fisher Scientific), and Vaseline, and the sandwich was transferred to a slide placed on a Peltier device on the microscope stage. Animals were imaged within 3 min of being removed from their cultivation temperature.

Animals were subjected to linear temperature ramps rising at 0.05°C/sec unless noted otherwise, via temperature-regulated feedback using LabView (National Instruments) and a T-type thermocouple (McShane Inc.). The slope of the temperature stimulus was selected to align with the temperature changes experienced by animals navigating the short thermal gradient. Based on average worm forward velocity of ∼0.15 mm/sec, animals are expected to experience temperature differences of 0.01°C/sec on the short thermal gradient. Since we observed few if any temperature responses when using a linear temperature ramp of 0.01°C/sec, we elected to use ramps rising at the rate of 0.05°C/sec to enable higher throughput analyses of responses. Individual animals were imaged for 4 mins at a rate of 2 Hz. Images were captured using a Zeiss 40X air objective (NA 0.9) or a Zeiss 10X air objective (NA 0.3) (for AFD and AIY imaging) on a Zeiss Axioskop2 Plus microscope, using a Hamamatsu Orca digital camera (Hamamatsu), and MetaMorph software (Molecular Devices). Data were analyzed using custom scripts in MATLAB (Mathworks) (Takeishi et al., 2016).

Each calcium trace was defined as the percent change in the relative fluorescence of the neuron from its baseline fluorescence level (average fluorescence of first 10 frames of each image) following background subtraction for all neurons with the exception of AFD. A fluorescence change of >10% in each neuron was considered a response, and the duration of calcium events was calculated as the sum of all events in each animal. Baseline fluorescence was set to zero to offset fluorescence change caused by photobleaching or movement artifacts. Calcium transients were imaged in the soma of AFD, AWC, AIB, and in the neurites of AIY, AIA and AIZ. *T\**_*AFD*_ was calculated as described previously (Takeishi et al., 2016).

### Quantification of *ins-1*p::GFP::PEST fluorescence

Young adult animals cultivated overnight at 20°C were well-fed throughout, or starved by placing them on unseeded NGM agar plates and re-cultivating at 20°C for 3h prior to imaging. Animals were moved onto 2% agarose pads on a glass slide, anesthetized with 1.5 µl of 25 mM azide, and were covered with a cover glass prior to imaging. Images were taken using an AX70 fluorescence microscope (Olympus) with an Olympus 20X UPlanSApo lens (NA 0.75). The ROI in the anterior intestine was outlined within 48 µm (75 pixels) from the posterior end of the pharynx and mean pixel intensity was calculated following background subtraction using ImageJ (NIH). Confocal microscope images were obtained using a FV3000 microscope (Olympus) with an Olympus 40X UPFLN lens (NA 0.75), and were exported as hyperstack .tif files.

### Statistical analyses

Excel (Microsoft) and GraphPad Prism version 8.0.0 (www.graphpad.com) were used to generate all histograms, bar graphs, and line-plots of cumulative distribution fractions. For statistical analysis of thermotaxis bias and chemotaxis, Student’s t-test was performed between fed and starved of each genotype and/or treatment. One-way ANOVA followed by Tukey’s multiple comparison was performed for Figures 1C and S1A using GraphPad Prism version 8.0.0. Wald F- and t-statistic analyses were performed using R (https://www.R-project.org/) and RStudio (www.rstudio.com) using the ‘emmeans’ package. To compare the distributions of reorientation direction following turns, the Mardia-Watson-Wheeler non-parametric test for circular data was performed using R and RStudio. All statistical analyses of imaging data were performed in MATLAB. The Kolmogorov-Smirnov test was performed to compare cumulative distribution fractions. The Mann-Whitney U test was used to compare the average duration of responses per neuron.

## Acknowledgements

We are grateful to Cori Bargmann, Steven Flavell, Oliver Hobert, Shawn Xu, Yun Zhang, and Joshua Hawk and Daniel Colon-Ramos for generously sharing reagents, the *Caenorhabditis* Genetics Center and the National BioResource Project (Japan) for strains, Masami Shima and the RIKEN CBS-Olympus Collaboration Center for experimental assistance, and Mike O’Donnell for assistance with statistics. We thank members of the Sengupta lab, Cori Bargmann, Miriam Goodman, Andrew Gordus and Yun Zhang for critical comments on the manuscript. This work was partly supported by the Japan Society for the Promotion of Science (H28-1058 – A.T.), and the NIH (T32 NS007292 and F32 NS112453 – N.H.; R35 GM122463 – P.S).

## Notes

### Competing Interest Statement

The authors have declared no competing interest.

