## Supplemental figures and tables for "Feeding state functionally reconfigures a sensory circuit to drive thermosensory behavioral plasticity"

### Supplemental Information

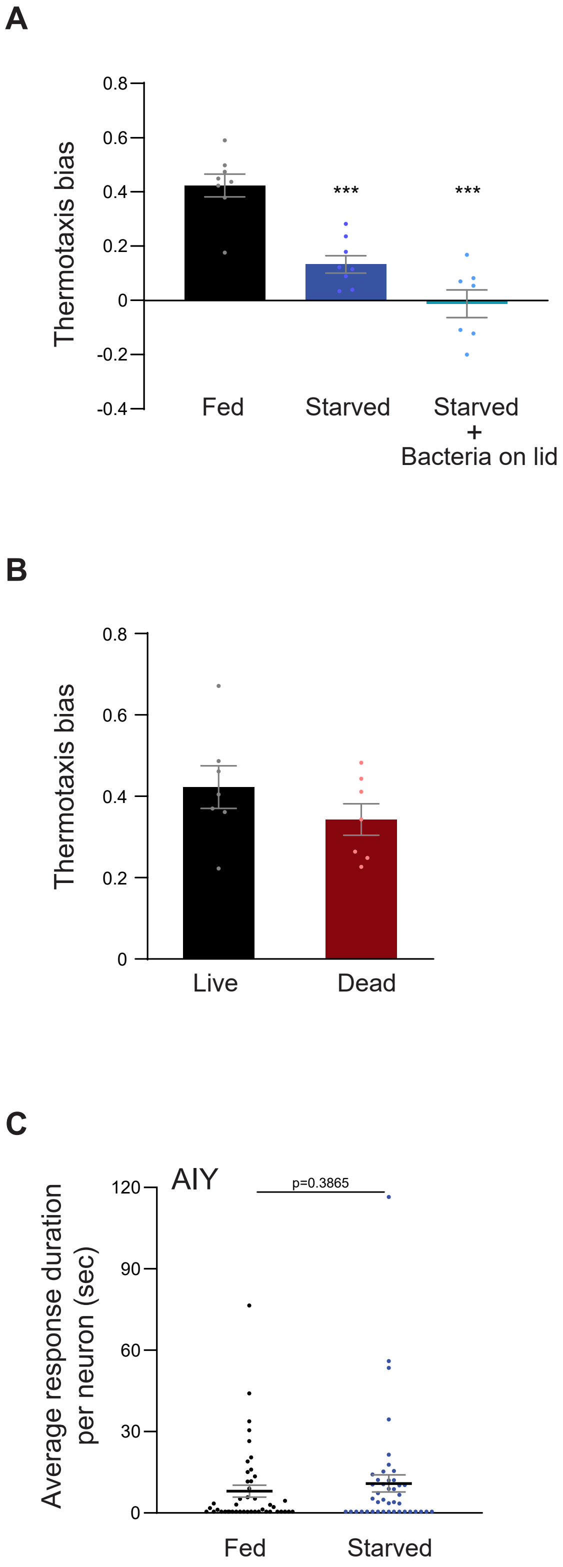


**Figure S1 related to Figure 1. Negative thermotaxis is modulated by internal feeding state.**

**A-B)** Mean thermotaxis bias of animals subjected to the indicated conditions. Animals were starved for 3h with or without exposure to OP50 bacteria present on the plate lids in **A**. Animals were fed with antibiotic-killed bacteria in **B** for 3h prior to the assay. Each dot represents the thermotaxis bias of a biologically independent assay comprised of 15 animals. Errors are SEM. *** indicates different from fed at p<0.001 (A: ANOVA with Tukey's multiple comparison test, B: Student's t-test).

**C)** Average duration of individual response events per AIY neuron (dots) in fed and starved animals, derived from data shown in Figure 1F. Means are indicated by horizontal black lines; errors are SEM. p-value was obtained using the Mann-Whitney U test.

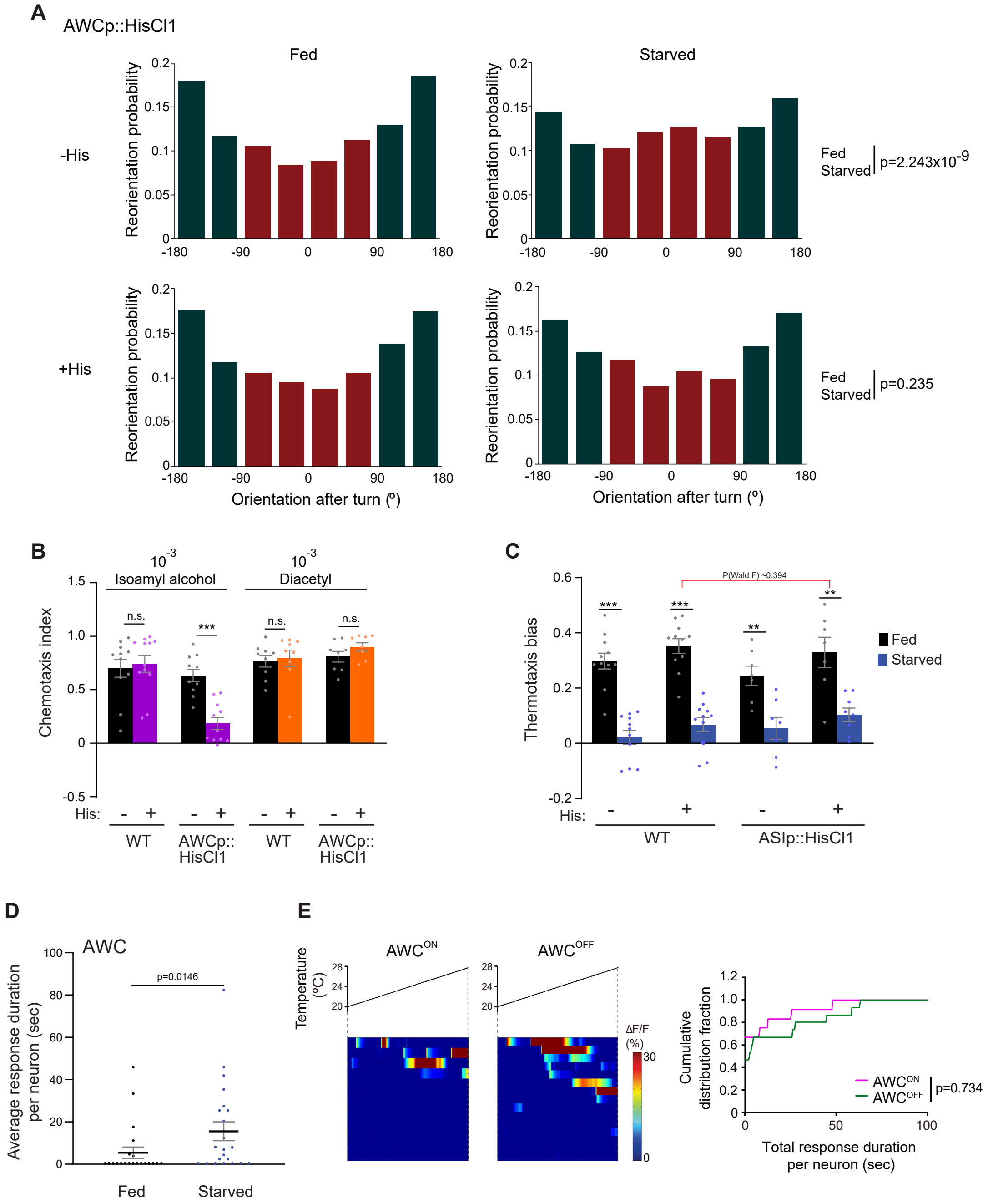


**Figure S2 related to Figure 2. Inhibition of AWC temperature responses restores negative thermotaxis behavior in starved animals.**

**A)** Histograms of movement orientation following a turn in animals expressing HisCl1 in AWC in the absence or presence of histamine on the assay plate. Data are from Figure 2A. Tracks from 8 assays of 15 animals each were categorized into bins of 45º. Red and green bars indicate orientation toward the warmer/orthogonal or cooler side, respectively. p-values were derived using the Mardia-Watson-Wheeler non-parametric test for circular data.

**B)** Mean chemotaxis index of animals of the indicated genotypes to a point source of the odorants isoamyl alcohol or diacetyl. Chemotaxis index is defined as [(number of animals at the odorant) – (number of animals at the control)/ total number of animals]. Assays were performed in the absence or presence of 10 mM histamine on the assay plate as indicated. Each dot represents a biologically independent assay of ~100 animals each. Errors are SEM. *** indicates different from fed (Student's t-test); n.s. – not significant.

**C)** Mean thermotaxis bias of fed and starved wild-type and transgenic animals expressing HisCl1 in ASI under the *srg-47* promoter in the presence or absence of 10 mM histamine on the assay plate. Each dot represents the thermotaxis bias of a biologically independent assay comprised of 15 animals. Errors are SEM. *** and ** indicate different from fed at each condition at $p<0.001$ and $p<0.01$, respectively (Student's t-test). P-value in red indicates Wald F-statistic from linear regression analysis for the effect of the indicated genotype on the magnitude of the feeding state effect. Wild-type data were interleaved with experimental data in Figure 2A, and Figure 3F-G, and are repeated.

**D)** Average duration of individual response events per AWC neuron (dots) in fed and starved animals, derived from data shown in Figure 2C. Means are indicated by horizontal black lines; errors are SEM. p-value was obtained using the Mann-Whitney U test.

**E)** (Left) Intracellular calcium dynamics in $AWC^{ON}$ and $AWC^{OFF}$ neurons in fed animals expressing GCaMP3 in response to a linear rising temperature stimulus (black lines) at 0.05ºC/sec. $AWC^{ON}$ and $AWC^{OFF}$ neurons were identified via the expression of *srsx-3*p*::mScarlet* in $AWC^{OFF}$. Each row in the heatmaps displays responses from a single neuron ordered by the time of the first response; n = 12 ($AWC^{ON}$) and 15 ($AWC^{OFF}$). (Right) Cumulative distribution fraction plots of total duration of calcium responses per $AWC^{ON}$ and $AWC^{OFF}$ neuron calculated from data shown at left. Distributions were compared using the Kolmogorov-Smirnov test.

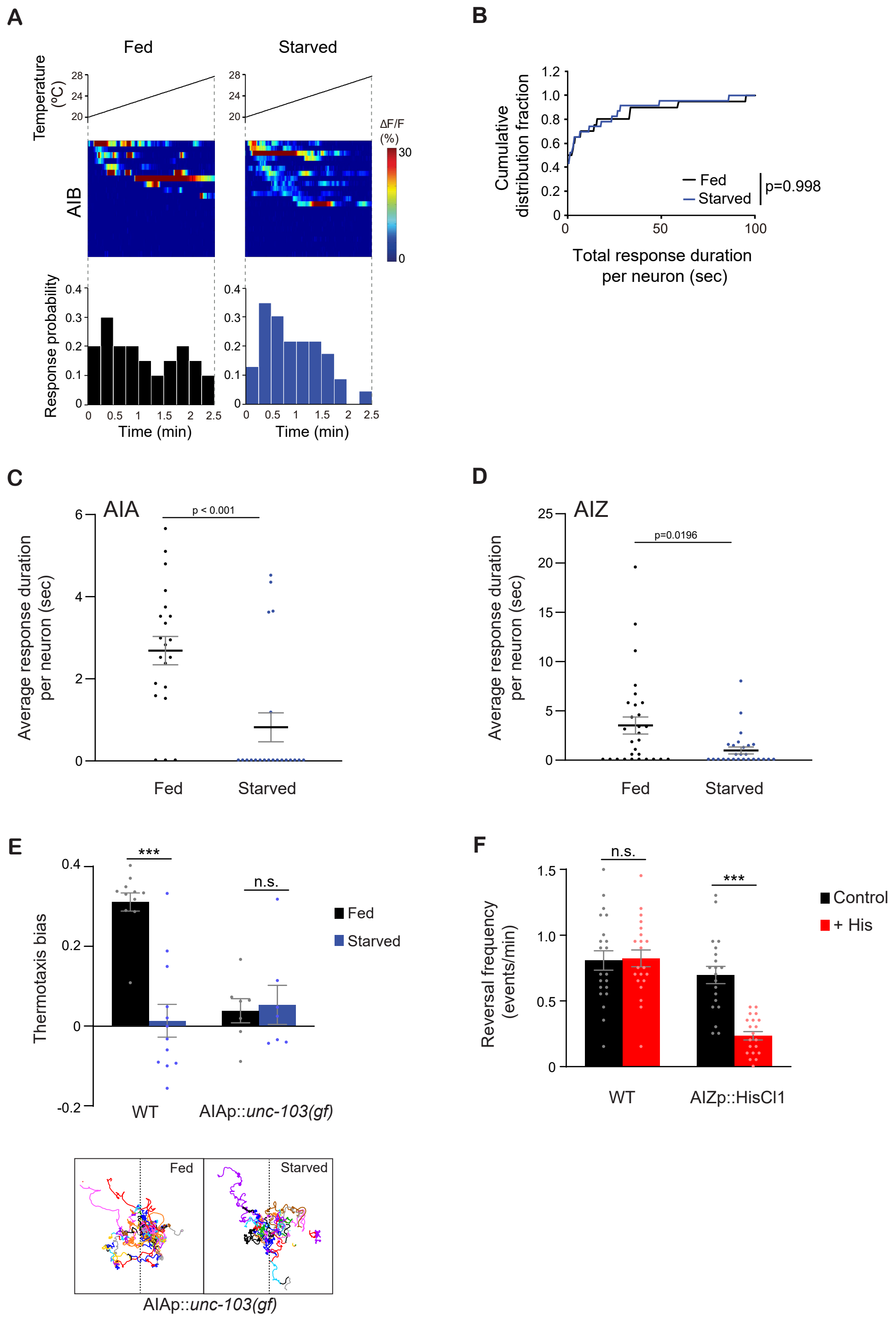


**Figure S3 related to Figure 3. Temperature responses in AIB are unaffected by starvation.**

**A)** Intracellular calcium dynamics in AIB expressing GCaMP3 in response to a linear rising temperature stimulus (black lines) at 0.05ºC/sec. Each row in the heatmaps displays responses from a single neuron ordered by the time of the first response; n = 20 (fed) and 23 (starved). (Bottom) Each bar in the histograms represents the percentage of neurons from animals of the indicated genotypes responding during 15s bins.

**B)** Cumulative distribution fraction plots of total duration of calcium responses per AIB neuron calculated from data shown in **A**. Distributions were compared using the Kolmogorov-Smirnov test.

**C-D)** Average duration of individual response events per AIA (**C**) and AIZ (**D**) neuron (dots) in fed and starved animals, derived from data shown in Figures 3B and 3D, respectively. Means are indicated by horizontal black lines; errors are SEM. p-values were obtained using the Mann-Whitney U test.

**E)** Mean thermotaxis bias of fed and starved wild-type and transgenic animals of the indicated genotypes (Table S1). Each dot represents the thermotaxis bias of a biologically independent assay comprised of 15 animals. Errors are SEM. *** indicates different from fed at p<0.001 (Student's t-test). n.s. – not significant. Wild-type data were interleaved with experimental data in Figures 3I, S4C, and S4F, and are repeated. Traces below show trajectories of fed and starved animals expressing *unc-103(gf)* in AIA; dashed lines indicate the starting temperature of 25.5ºC on the thermal gradient. Individual worm trajectories are color-coded. Since trajectories are terminated by omega turns or collisions, trajectories of the same color may not represent the movement of a single animal throughout the assay.

**F)** Mean reversal frequency of fed and starved wild-type and transgenic animals expressing HisCl1 in AIZ in the presence or absence of 10 mM histamine on an isothermal plate at 20ºC. Each dot represents the reversal frequency of an individual animal over the assay period. n = 20. Errors are SEM. *** indicates different from fed (Student's t-test); n.s. – not significant.

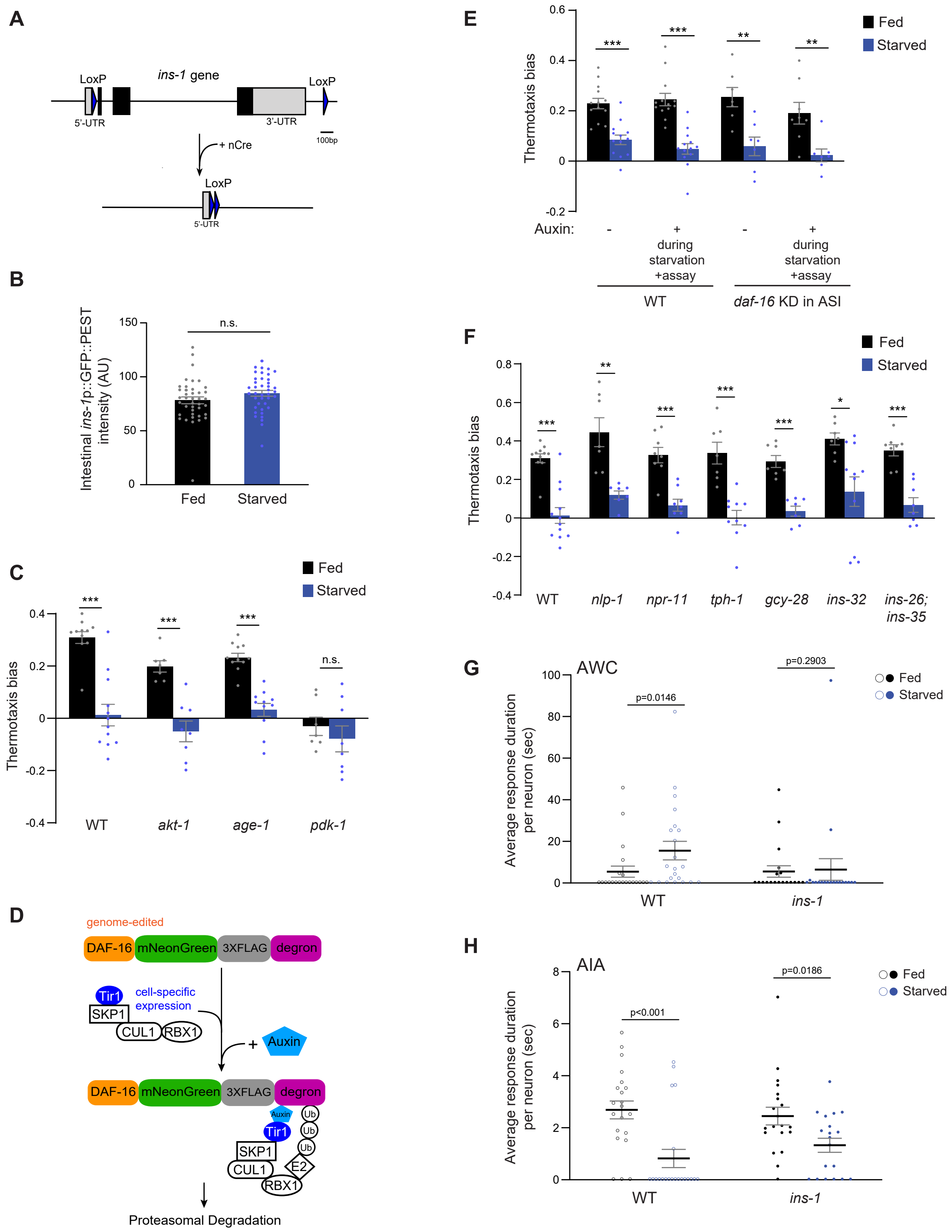


**Figure S4 related to Figure 4. Insulin signaling mediates starvation-dependent thermotaxis behavioral plasticity.**

**A)** Schematic of Cre-Lox-mediated recombination strategy for cell-specific knockout of *ins-1*. *loxP* sequences were inserted flanking the *ins-1* genomic sequence via gene editing. nCre was expressed under cell-specific promoters (Table S1).

**B**) Average pixel intensity of *ins-1*p::GFP::PEST fluorescence in the anterior intestine of well-fed and starved (3h) animals. Each dot is the average fluorescence intensity in a single animal. Errors are SEM. n.s. – not significant.

**C, E, F)** Mean thermotaxis bias of fed and starved animals of the indicated genotypes (see Table S1 for alleles used). Each dot represents the thermotaxis bias of a biologically independent assay comprised of 15 animals. Errors are SEM. DAF-16 was depleted in ASI via auxin-induced degradation of a degron-tagged *daf-16* allele and ASI-specific expression of TIR1 under the *srg-47* promoter **(E)**. Data from two independent transgenic strains are combined (**E**). Auxin was added during starvation and to the assay plate. ***, ** and * indicate different from fed at p<0.001, p<0.01, and p<0.05, respectively (Student's t-test). n.s. – not significant. Wild-type data in **C** and **F** were interleaved with experimental data in Figures 3I and S3E, and are repeated. Wild-type data in **E** were interleaved with experimental data in Figure 4E, and are repeated.

**D)** Schematic of auxin-induced degradation strategy. The *daf-16* gene edited allele was a kind gift of Oliver Hobert (Columbia University) (Aghayeva et al., 2020). TIR1-encoding sequences were expressed under cell-specific promoters (Table S2).

**G, H)** Average duration of individual response events per AWC (**G**) and AIA (**H**) neuron (dots) in fed and starved wild-type and *ins-1* mutants. Wild-type data are repeated (open circles) from Figures S2D (**G**) and S3C (**H**). *ins-1* data are derived from Figures 4F and 4G. Means are indicated by horizontal black lines; errors are SEM. p-values were obtained using the Mann-Whitney U test.

**Table S1 related to all Figures.** Strains used in this work.

| **Strain** | **Genotype** | **Source/parent strains** | **Relevant Figures** |
|---|---|---|---|
| WT | N2 (Bristol) | CGC | |
| DCR3055 | *wyIs629*[*gcy-8*p::*GCaMP6s* + *gcy-8*p::*mCherry* + *unc-122*p::*gfp*] | (Hawk et al., 2018) | 1E |
| DCR3056 | *olaIs17*[*mod-1*p::*GCaMP6s* + *ttx-3*p::*mCherry* + *unc-122*p::*dsRed*] | (Hawk et al., 2018) | 1F, 1G, S1C |
| PY12205 | *oyEx657*[*odr-1*p::*HisCl1*::*SL2*::*mCherry* + *unc-122*p::*gfp*] | This paper | 2A, S2A, S2B |
| SWF147 | *flvEx76*[*ceh-36(prom3)*p::*GtACR2*::*T2A*::*gfp* + *myo-3*p::*mCherry*] | Gift from Steven Flavell | 2B |
| PY11012 | *oyEx658*[*odr-1*p::*GCaMP3* + *srsx-3*p::*mScarlet* + *unc-122*p::*dsRed*] | This paper | 2C, 2D, 4F, 4H, S2D, S2E, S4G |
| CX15257 | *kyEx5128*[*gcy-28d*p::*GCaMP5A* + *unc-122*p::*dsRed*] | (Larsch et al., 2015) | 3B, 3C, 4G, 4H, S3C, S4H |
| ZC2467 | *yxEx1364*[*ser-2(2)*p::*FRT*::*GCaMP6s* + *odr-2b(3a)*p::*nFLP* + *unc-122*p::*dsRed*] | This paper | 3D, 3E, S3D |
| CX15261 | *kyIs617*[*gcy-28d*p::*HisCl1*::*SL2*::*gfp* + *myo-3*p::*mCherry*] | (Cho et al., 2016) | 3F |
| PY12206 | *oyEx659*[*ser-2(2)*p::*FRT*::*HisCl1*::*SL2*::*mCherry* + *odr-2d(3a)*p::*nFLP* + *unc-122*p::*gfp*] | This paper | 3G, S3F |
| CX17432 | *kyEx6105*[*ins-1(s)*p::*Chrimson*::*SL2*::*mCherry* + *elt-2*p::*mCherry*]; *kyEx5128*[*gcy-28d*p::*GCaMP5A* + *unc-122*p::*dsRed*] | (Lopez-Cruz et al., 2019) | 3H |
| CX17524 | *kySi77*[*FRT* before *eat-4* start codon] *kySi76*[*let-858UTR*::*FRT*::*mCherry* after *eat-4* stop codon] III; *kyEx6142*[*odr-1*p::*nFLP*] | López-Cruz et al., 2019 | 3I |
| PY12200 | *ins-1*(*oy158*)[*loxP*::*ins-1*::*loxP*] | This paper | 4A |
| PY12201 | *ins-1(oy158)*; *oyEx660*[*ins-1*p::*nCre*::*SL2*::*gfp* +*unc-122*p::*dsRed]* | Injected into PY12200 | 4A |
| PY12202 | *ins-1(oy158)*; *oyEx661*[*ifb-2*p::*nCre*::*SL2*::*gfp* +*unc-122*p::*dsRed*] | Injected into PY12200 | 4A |
| PY12203 | *ins-1(oy158); oyEx662*[*gcy-28d*p::*nCre*::*SL2*::*gfp* +*unc-122*p::*dsRed*] | Injected into PY12200 | 4A |
| PY12204 | *ins-1(oy158); oyEx663*[*odr-2(3a)*p::*nCre*::*SL2*::*gfp* +*unc-122*p::*dsRed*] | Injected into PY12200 | 4A |
| PY11612 | *oyIs90*[*ins-1*p::*gfp*::*PEST* + *unc-122*p::*dsRed*] | This paper | 4B, S4B |
| CX7155 | *ins-1*(*nr2091*) | (Pierce et al., 2001) | 4D |
| DR1572 | *daf-2*(*e1368*) | CGC | 4D |
| DR26 | *daf-16*(*m26*) | CGC | 4D |
| RB712 | *daf-18*(*ok480*) | CGC | 4D |

| | | | |
|---|---|---|---|
| PY12207 | OH14125[*ot853*[*daf-16*p::linker::*mNeonGreen*::*3xflag*::*AID*]]; *oyEx664*[*ceh-36prom2_del1ASE*::*TiR1*::*SL2*::*mTurquiose2* + *unc-122p*::*dsRed* + *N2 gDNA*] | Injected into OH14125 | 4E |
| PY11614 | *ins-1*(*nr2091*); *oyEx658*[*odr-1*p::*GCaMP3* + *str-2*p::*mScarlet* + *unc-122*p::*dsRed*] | CX7155, PY11012 | 4F, 4H, S4G |
| PY11615 | *ins-1*(*nr2091*); *kyEx5128*[*gcy-28d*p::*GCaMP5A* + *unc-122*p::*dsRed*] | CX7155, CX15257 | 4G, 4H, S4H |
| PY12208 | *oyEx665*[*srg-47*p::*HisCl1*::*SL2*::*mCherry* + *unc-122*p::*gfp*] | This paper | S2C |
| CX13440 | *kyEx4018*[*inx-1*p::*GCaMP3* + *unc-122*p::*dsRed*] | (Gordus et al., 2015) | S3A, S3B |
| CX14599 | *kyEx4747*[*gcy-28d*p::*unc-103(gf)*::*SL2*::*mCherry* +*elt-2*p::*mCherry]* | (Cho et al., 2016) | S3E |
| RB759 | *akt-1*(*ok525*) | CGC | S4C |
| TJ1053 | *age-1*(*hx546*) | CGC | S4C |
| JT9609 | *pdk-1*(*sa680*) | CGC | S4C |
| PY11616 (line 1) | OH14125[*ot853*[*daf-16*p::linker::*mNeonGreen*::*3xFlag*::*AID*]; *oyEx666*[*srg-47*p::*TiR1*::*SL2*::*mTurquiose2* + *unc-122*p*::dsRed + N2 gDNA]* | Injected into OH14125 | S4E |
| PY11617 (line 2) | OH14125[*ot853*[*daf-16*p::linker::*mNeonGreen*::*3xFlag*::*AID*]; *oyEx667*[*srg-47*p::*TiR1*::*SL2*::*mTurquiose2* + *unc-122*p*::dsRed + N2 gDNA]* | Injected into OH14125 | S4E |
| RB1341 | *nlp-1*(*ok1470*) | CGC | S4F |
| RB799 | *npr-11*(*ok594*) | CGC | S4F |
| MT15434 | *tph-1*(*mg280*) | CGC | S4F |
| FX02411 | *gcy-28*(*tm2411*) | NBRP | S4F |
| PY10808 | *ins-32*(*tm6109*) | NBRP | S4F |
| PY10712 | *ins-26*(*tm1983*); *ins-35*(*ok3297*) | PY10708, RB2412 | S4F |

**Table S2 related to all Figures.** Plasmids used in this work.

| Plasmid | Sequences | Promoter length | Source |
|---|---|---|---|
| PSAB1201 | *odr-1*p::*GCaMP3* | 1.0 kb | This paper |
| PSAB1202 | *srg-47*p::*GCaMP3* | 650 bp | This paper |
| PSAB1203 | *srsx-3*p::*mScarlet* | 1.3 kb | This paper |
| PSAB1204 | *odr-1*p::*HisCl1*::*SL2*::*mCherry* | 1.0 kb | This paper |
| PSAB1205 | *srg-47*p::*HisCl1*::*SL2*::*mCherry* | 650 bp | This paper |
| PSAB1206 | *ser-2(2)*p::*FRT*::*HisCl1*:: *SL2*::*mCherry* | 4.7 kb | This paper |
| WY016 | *ser-2(2)*p*::FRT::STOP::FRT::GCaMP6s* | 4.7 kb | This paper |
| PSAB1207 | *ins-1*p::*nCre*::*SL2*::*gfp* | 4.2 kb | This paper |
| PSAB1208 | *gcy-28d*p::*nCre*::*SL2*::*gfp* | 2.8 kb | This paper |
| PSAB1209 | *odr-2b(3a)*p::*nCre*::*SL2*::*gfp* | 441 bp | This paper |
| PSAB1210 | *ifb-2*p::*nCre*::*SL2*::*gfp* | 3.0 kb | This paper |
| PSAB1211 | *ins-1*p:*:gfp*::*PEST* | 4.2 kb | This paper |
| PSAB1212 | *srg-47*p::*Tir1*::*mTurquoise2* | 650 bp | This paper |
| pWY046 | *odr-2b(3a)*p::*nFlp* | 441 bp | This paper |
| pUA118 | *ceh-36prom2_del1ASE*p::*TiR1*:: *SL2*::*mTurquoise2* | 1.6 kb | Gift from Oliver Hobert |
